# Physiological and head motion signatures in static and time-varying functional connectivity and their subject discriminability

**DOI:** 10.1101/2020.02.04.934554

**Authors:** Alba Xifra-Porxas, Michalis Kassinopoulos, Georgios D. Mitsis

## Abstract

Human brain connectivity yields significant potential as a noninvasive biomarker. Several studies have used fMRI-based connectivity fingerprinting to characterize individual patterns of brain activity. However, it is not clear whether these patterns mainly reflect neural activity or the effect of physiological and motion processes. To answer this question, we capitalize on a large data sample from the Human Connectome Project and rigorously investigate the contribution of the aforementioned processes on functional connectivity (FC) and time-varying FC, as well as their contribution to subject identifiability. We find that head motion, as well as heart rate and breathing fluctuations, induce artifactual connectivity within distinct resting-state networks and that they correlate with recurrent patterns in time-varying FC. Even though the spatiotemporal signatures of these processes yield above-chance levels in subject identifiability, removing their effects at the preprocessing stage improves identifiability, suggesting a neural component underpinning the inter-individual differences in connectivity.

## 1 Introduction

Functional magnetic resonance imaging (fMRI) is based on the blood-oxygenation-level-dependent (BOLD) contrast mechanism (Ogawa et al., 1990), and is widely viewed as the gold standard for studying brain function because of its high spatial resolution and non-invasive nature. The BOLD signal exhibits low frequency (~0.01-0.15 Hz) fluctuations that are synchronized across different regions of the brain, a phenomenon known as functional connectivity (FC). FC has been observed even in the absence of any explicit stimulus or task, giving rise to the so-called resting-state networks (RSNs) (Biswal et al., 1995; Fox and Raichle, 2007; Smith et al., 2009). Initially, FC was viewed as a stationary phenomenon (static FC) and was commonly measured as the correlation between brain regions over an entire scan. However, several researchers challenged this assumption (Chang and Glover, 2010; Sakoglu et al., 2010), and recent studies have been focusing on FC dynamics, quantified over shorter time scales than the scan duration (time-varying FC) (Hutchison et al., 2013; Lurie et al., 2019).

Although the neurophysiological basis of resting-state FC measured with fMRI is not yet fully understood, many studies have provided evidence to support its neuronal origin. For instance, in animal models, a strong association between spontaneous BOLD fluctuations and neural activity, in particular band-limited local field potentials and firing rates, has been reported (Logothetis et al., 2001; Schölvinck et al., 2010; Shmuel and Leopold, 2008; Thompson et al., 2013b). Furthermore, a recent study suggested a close correspondence between windowed FC calculated from simultaneously recorded hemodynamic signals and calcium transients (Matsui et al., 2018). In human studies, direct measurements of macroscale neural activity have revealed a spatial correlation structure similar to that of spontaneous BOLD fluctuations (Brookes et al., 2011; Hacker et al., 2017; He et al., 2008; Hipp et al., 2012; Kucyi et al., 2018), even during transient (50-200 ms) events (A. P. Baker et al., 2014; Hunyadi et al., 2019; Vidaurre et al., 2018). Therefore, it is widely assumed that resting-state FC measured using BOLD fMRI reflects spontaneous co-fluctuations of the underlying neuronal networks.

However, BOLD signals rely on changes in local cerebral blood flow (CBF) to infer underlying changes in neuronal activity, and according to a recent study, at least 50% of the spontaneous hemodynamic signal is unrelated to ongoing neural activity (Winder et al., 2017). For instance, systemic physiological functions can induce variations in global and local CBF, which in turn result in BOLD signal fluctuations. In particular, low frequency variations in breathing activity (Birn et al., 2008b, 2006; Power et al., 2017), arterial blood pressure (Whittaker et al., 2019), arterial CO_2_ concentration (Prokopiou et al., 2019; Wise et al., 2004), and heart rate (Chang et al., 2009; Shmueli et al., 2007) are known to account for a considerable fraction of variance of the BOLD signal, presumably through changes in CBF. In addition, the BOLD signal intensity is distorted by high-frequency physiological fluctuations, such as cardiovascular pulsation and breathing, through displacement of the brain tissues and perturbations of the B0 magnetic field (Dagli et al., 1999; Glover et al., 2000). Further, head motion is well-known to have a substantial impact on fMRI through partial volume, magnetic inhomogeneity and spin-history effects (Friston et al., 1996; Power et al., 2012). These non-neuronal factors may introduce common variance components in signals recorded from different brain regions and subsequently induce spurious correlations between these areas (Chen et al., 2020). Therefore, to account for motion-related and physiological confounds, nuisance regressors are typically obtained using model-based and data-driven techniques, and regressed out from the fMRI data before further analysis (Caballero-Gaudes and Reynolds, 2017).

Static and time-varying resting-state FC have shown promise for providing concise descriptions of how the brain changes across the lifespan (Battaglia et al., 2017; Chan et al., 2014; Ferreira et al., 2016; Geerligs et al., 2015; Sala-Llonch et al., 2015; Xia et al., 2019), and to assay neural differences that are associated with disease (J. T. Baker et al., 2014; Chen et al., 2017; Damaraju et al., 2014; Demirtaş et al., 2016; Drysdale et al., 2017; Du et al., 2016; Gratton et al., 2019; Hahamy et al., 2015; Mash et al., 2019; Morgan et al., 2017; Xia et al., 2018). However, recent studies assessing the performance of a large range of preprocessing strategies found that there is always a trade-off between adequately removing confounds from fMRI data and preserving the signal of interest (Ciric et al., 2017; Kassinopoulos and Mitsis, 2019a; Parkes et al., 2018). Importantly, these studies found that widely used techniques for the preprocessing of fMRI data may not efficiently remove physiological and motion artifacts. The latter raises a concern, as it is still not clear how nuisance fluctuations may impact the outcome of FC studies.

Several studies have examined whether physiological fluctuations across the brain could give rise to structured spatial patterns that resemble common RSNs, based on the evidence that vascular responses following systemic changes are spatially heterogenous (Chang et al., 2009; Pinto et al., 2017), or account for the observed time-varying interactions between RSNs. For instance, (Bright and Murphy, 2015) applied independent component analysis (ICA) to the fraction of the fMRI data explained by nuisance regressors related to head motion and physiological variability and revealed a characteristic network structure similar to previously reported RSNs. Similarly, (Tong et al., 2013; Tong and Frederick, 2014) found significant contributions of systemic fluctuations on ICA time courses related to the visual, sensorimotor and auditory networks. Recently, (Chen et al., 2020) generated BOLD data containing only slow respiratory-related dynamics and showed that respiratory variation can give rise to apparent neurally-related connectivity patterns. Further, recent investigations have shown that physiological confounds can modulate time-varying FC measures (Chang et al., 2013; Nalci et al., 2019; Nikolaou et al., 2016). These results suggest that changes in brain physiology, breathing patterns, heart rhythms and head motion across sessions, within-subject or across populations, may introduce artifactual inter-individual and group-related differences in FC independent of any underlying differences in neural activity. For instance, cardiac autonomic dysregulation has been associated to a variety of psychiatric disorders (Alvares et al., 2016; Benjamin et al., 2020), which could in principle lead to group differences in connectivity patterns between patients and controls if the effects of heart rate are not accounted for. Therefore, the disentanglement of the neural and physiological correlates of resting-state FC is crucial for maximizing its clinical impact.

While the previous findings provide evidence for the dual-nature of RSNs in both static and time-varying scenarios, only specific physiological processes and/or particular brain networks were evaluated in each of the aforementioned studies. A more holistic assessment of the impact of these non-neural processes on FC measures is needed to better understand whether and how systemic fluctuations, as well as head or breathing motion affect inter-individual and group differences. Importantly, the wide range of possible preprocessing strategies needs to be reassessed taking into consideration the effects of several non-neural processes on FC measures rather than accounting only for the effects of a specific process (e.g. head motion).

The varying efficiency of different preprocessing pipelines with respect to removing the effect of physiological fluctuations and motion has also implications for studies investigating properties of FC at the individual level. Recent studies have shown that connectivity profiles vary substantially between individuals, acting as an identifying fingerprint (Finn et al., 2015; Mira-Dominguez et al., 2014) that is stable over long periods of time (Horien et al., 2019). However, the high subject discriminability of connectivity profiles may arise partly as a result of physiological processes (Batchvarov et al., 2002; Golestani et al., 2015; Malik et al., 2008; Pinna et al., 2007; Pitzalis et al., 1996; Power et al., 2020; Reland et al., 2005) and head motion (van Dijk et al., 2012; Zeng et al., 2014) being highly subject-specific. Evidence supporting this hypothesis comes from studies showing that the mean of intraclass correlation values of functional connections, associated to test-retest reliability across sessions, is reduced when a relatively aggressive pipeline is used (Birn et al., 2014; Kassinopoulos and Mitsis, 2019a; Parkes et al., 2018), suggesting that artifacts exhibit high subject specificity. However, the relation between subject discrimination and inter-individual differences in physiological processes and head motion has yet to be addressed.

In the present work, we capitalize on 3T resting-state fMRI data from the Human Connectome Project (HCP) to uncover the whole-brain connectome profiles of systemic low-frequency oscillations (SLFOs) associated with heart rate and breathing patterns, cardiac pulsatility, breathing motion and head motion on estimates of static and time-varying FC. To quantify the contributions of physiological processes and head motion on FC, we employ model-based techniques with externally recorded physiological measurements, and subsequently generate nuisance datasets that only contain non-neural fluctuations. Using these datasets, we provide a comprehensive examination of the regional variability of the impact of the considered nuisance processes on the BOLD signal, as well as an investigation of the group consistency and inter-individual differences of their characteristic signatures on FC. We further evaluate several fMRI preprocessing strategies to assess the extent to which different techniques remove the physiological and motion FC signatures from the fMRI data. Finally, we investigate the potential effect of physiological processes and head motion on individual discriminability in the context of connectome fingerprinting. Using the proposed approach, we show that SLFOs and head motion have a larger impact on FC measures compared to breathing motion and cardiac pulsatility, and we highlight the functional connections that are more prone to exhibiting biases. Furthermore, our findings suggest that the recurrent whole-brain connectivity patterns observed in time-varying FC can be partly attributed to SLFOs and head motion. Finally, we show that connectome fingerprinting accuracies are higher when non-neural confounds are reduced, suggesting a neural component underpinning the individual nature of FC patterns.

The codes that were employed to carry out the analyses described in the present study are publicly available and can be found on github.com/axifra/Nuisance_signatures_FC.

## 2 Results

### 2.1 Contributions of nuisance processes on the BOLD signal

We examined regional differences in the influence of physiological processes and head motion on the BOLD signal. The physiological processes evaluated here were breathing motion, cardiac pulsatility, and SLFOs associated with changes in heart rate and breathing patterns (see Materials and methods – Nuisance processes evaluated). Scans with LR and RL phase encoding were examined separately as it has been suggested that breathing motion artifacts vary across scans with different phase encoding directions (Raj et al., 2001), and thus we aimed to examine whether other processes such as head motion demonstrate a similar dependence. The contributions of each nuisance process on BOLD signal fluctuations were quantified as the correlation between the nuisance fluctuations of the process in question, modelled using externally recorded physiological measurements, and the BOLD fluctuations “cleaned” of all other nuisance fluctuations (denoted as *r_nuis_* in the Materials and methods section – Isolation of nuisance fluctuations from fMRI data, see also **Figure 7**). We computed these contributions for each scan and then tested for the presence of consistent patterns across scans with the same phase encoding direction (significance testing using inter-subject surrogates, two-sample t-test, *p*<0.05, Bonferroni corrected).

The results showed distinct regional patterns for each of the nuisance processes. SLFOs mostly affected sensory regions, including the visual and somatosensory cortices (particularly of the face) (**Figure 1A**). Phase encoding was not found to modulate the magnitude of the SLFOs contributions on the BOLD signal (**Figure 1A**). Head motion exhibited the largest effect in the somatosensory and visual cortices (**Figure 1B**). Intriguingly, the effect in the visual cortex was highest in the right hemisphere for LR phase encoding, but highest in the left hemisphere for RL phase encoding (**Figure 1B**). Breathing motion effects were more pronounced in prefrontal, parietal and temporal brain regions (**Figure 1C**). Further, breathing motion had a much larger impact on the left hemisphere when the phase encoding was LR, whereas the reverse pattern was observed for RL phase encoding (**Figure 1C**). Cardiac pulsatility was highest in regions such as the visual and auditory cortices, as well as the insular cortex (**Figure 1D**).

**Figure 1.**
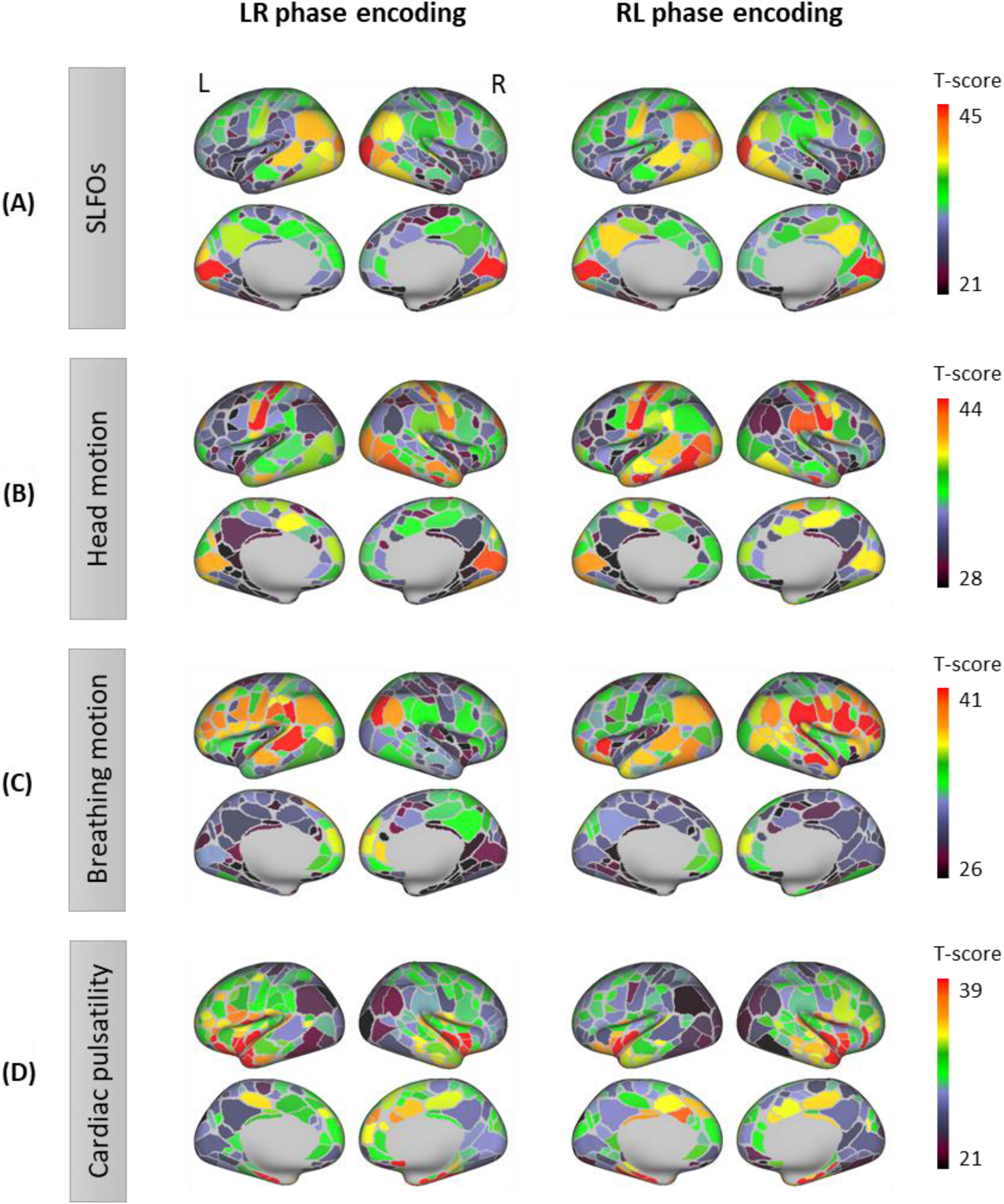
Contributions of nuisance processes on the resting-state BOLD signal. *T*-score maps of the correlation between each nuisance process and BOLD fMRI fluctuations (raw data) for **(A)** SLFOs, **(B)** head motion, **(C)** breathing motion, and **(D)** cardiac pulsatility, computed within each parcel of the Gordon atlas (two-sample *t*-test against the surrogate data, *p*<0.05, Bonferroni corrected). The *t*-tests were calculated for each phase encoding separately. The physiological fluctuations were obtained from simultaneous external recordings. These results illustrate the cortical regions most affected by each nuisance process.

### 2.2 Physiological and head motion signatures in static FC

To examine the effect of physiological fluctuations and head motion in static FC, we developed a framework that quantifies the extent to which functional connections are influenced by a nuisance process at the individual scan level. Briefly, synthetic datasets were generated for each scan based on the contributions of the examined nuisance processes within each ROI (see Materials and methods – Isolation of nuisance fluctuations from fMRI data). These datasets retained the variance explained by nuisance fluctuations and replaced the remaining variance (often considered as the “neural” variance) with uncorrelated random signals. This framework allowed us to compute FC matrices that illustrate the whole-brain connectome profiles arising from the nuisance processes of interest (see Materials and methods – Estimation of static and time-varying functional connectivity).

The group-averaged static FC matrices across all 1,568 scans revealed consistent whole-brain connectome patterns for SLFOs, head motion and breathing motion (**Figure 2A-C**). SLFO-based connectivity profiles exhibited strong positive correlations for all edges of the FC matrix, particularly for edges within the visual network, as well as between the visual network and the rest of the brain (**Figure 2A**). Head motion mainly influenced functional connections within the visual and sensorimotor networks, as well as edges within the DMN (**Figure 2B**). Note that even though areas in both the visual and sensorimotor networks were influenced by motion artifacts (**Figure 1B**), we did not observe strong correlations between the two aforementioned networks. This is not entirely surprising, as two brain areas may be associated with a different linear combination of head motion nuisance regressors and, thus, the correlation between the region-specific motion-induced fluctuations can in principle be around zero. Breathing motion exhibited an intriguing chess-like pattern, with both positive and negative correlations (**Figure 2C**, lower triangular matrix). Based on this observation, we subsequently reordered the ROIs with respect to their hemisphere, which revealed that positive correlations were mostly confined between ROIs of the same hemisphere, whereas correlations between hemispheres were close to zero or even negative (**Figure 2C**, upper triangular matrix). Even when scans with LR and RL phase encoding were averaged separately, both hemispheres exhibited increased within-hemisphere connectivity (**Supp. Fig. 1C**). Nonetheless, the connectome profile of breathing motion exhibited clear differences between phase encoding directions, whereas all other nuisance processes did not exhibit perceivable differences (**Supp. Fig. 1**). Finally, cardiac pulsatility did not exhibit a characteristic spatial pattern and the group-averaged correlation values were low, suggesting that it does not affect static FC in a systematic manner across subjects (**Figure 2D**).

**Figure 2.**
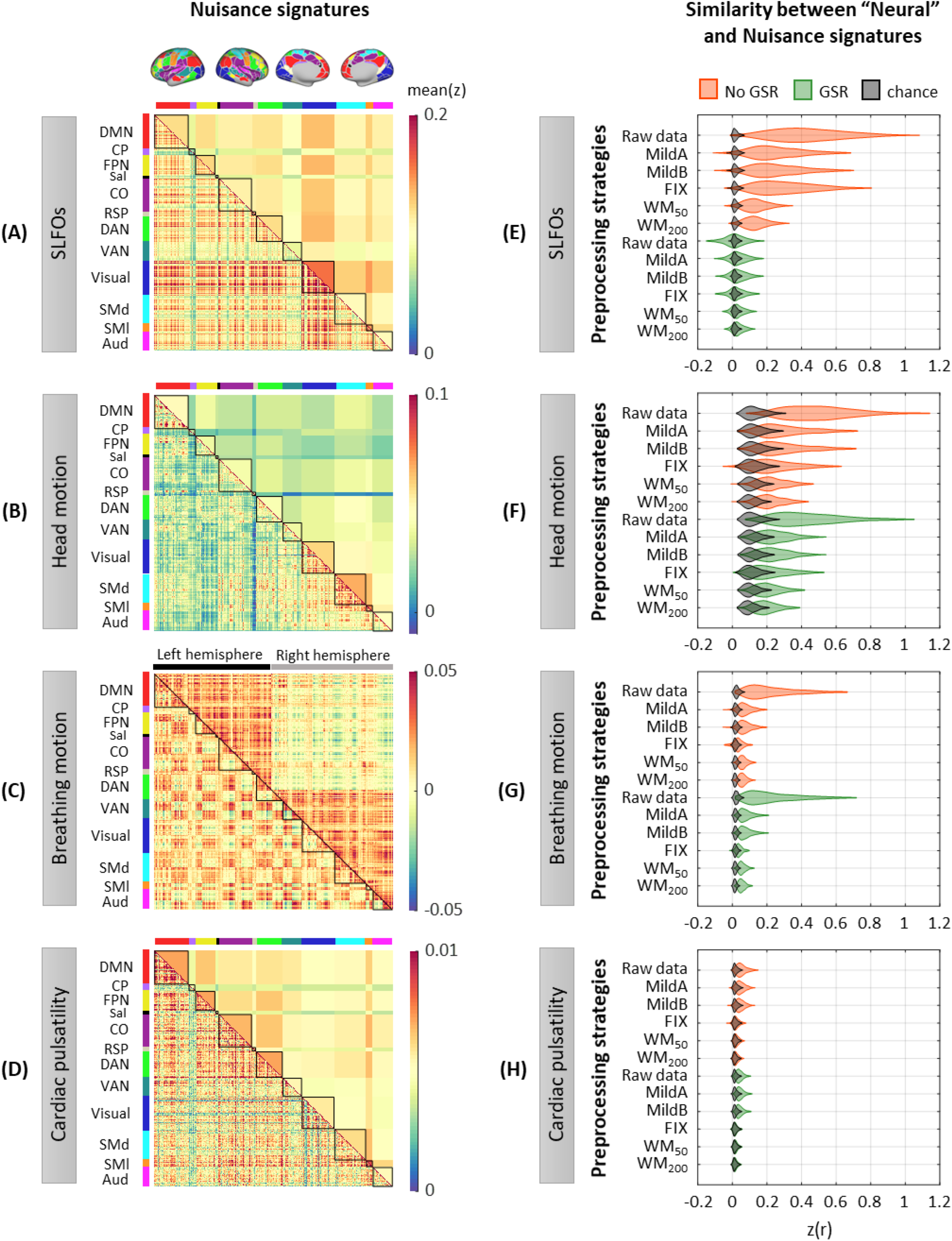
Whole-brain connectome patterns induced by nuisance processes and effect of preprocessing strategies. (**A-D**) Group averaged nuisance FC matrices across all 1,568 scans for (**A**) SLFOs, (**B**) head motion, (**C**) breathing motion, and (**D**) cardiac pulsatility. These results demonstrate that nuisance fluctuations induce heterogeneous whole-brain connectivity profiles which, if unaccounted for, can result in biased estimates functional connectivity. (**E-H)** Distribution of Pearson correlation coefficients across all 1,568 scans between the “neural” FC matrix for different preprocessing strategies and nuisance FC matrices associated to (**E**) SLFOs, (**F**) head motion, (**G**) breathing motion, and (**H**) cardiac pulsatility. Correlation values were Fisher z transformed. SLFOs, head motion and breathing motion were found to confound the FC matrices more severely (E-G). GSR effectively removed the effects of SLFOs, while more aggressive preprocessing pipelines mitigated the effects of head motion, breathing motion and cardiac pulsatility.

### 2.3 Capability of preprocessing strategies to remove the nuisance signatures on static FC

To examine the capability of various preprocessing strategies to reduce the effects introduced by physiological processes and head motion on static FC, we computed for each scan the similarity of the connectome profile that arises from a nuisance process with the connectome profile calculated from preprocessed fMRI data (considered as the “neural” profile). This similarity reflects the extent to which the “neural” connectome profile extracted after a specific denoising strategy is confounded by physiological and head motion artifacts. A distribution of the similarity values across scans, in this case Pearson’s correlation coefficients, is shown in **Figure 2E-H** for each preprocessing strategy and nuisance process. We found that SLFOs, head motion and breathing motion had the strongest influence on static FC, based on the similarity of their connectome profiles with the “neural” connectome profiles from the raw data (**Figure 2E-H**).

The signature induced by SLFOs remained after the MildA, MildB and FIX pipelines were applied, but was greatly reduced by the WM_50_ and WM_200_ strategies (**Figure 2E**). The observation that FIX, which is a rather aggressive preprocessing strategy, was unable to remove most of the SLFOs is consistent with recent studies showing that global artifactual fluctuations are still prominent after FIX denoising (Burgess et al., 2016; Glasser et al., 2018; Kassinopoulos and Mitsis, 2019b; Power et al., 2018, 2017). Notably, GSR seemed to be an effective technique for removing the physiological signature from SLFOs on static FC, albeit for some scans it appeared to introduce a negative correlation between the SLFOs and “neural” FC matrices (**Figure 2E**). This effect was greater when the global signal was computed across the whole brain in volumetric space, compared to across vertices in surface space (**Supp. Fig. 2A**). The effect of the signature related to head motion was reduced with more aggressive preprocessing strategies, but none of the examined approaches completely eradicated the head motion effects in static FC (**Figure 2F**). GSR slightly reduced the similarity between the head motion and “neural” connectome profiles. The signature induced by breathing motion was greatly reduced by all preprocessing strategies, and particularly by FIX denoising, which yielded almost chance level (**Figure 2G**). Still, none of the preprocessing strategies entirely eliminated the breathing motion signature in static FC. The confounds introduced by cardiac pulsatility were overall small and effectively removed by the FIX, WM_50_ and WM_200_ strategies (**Figure 2H**). GSR did not have any effect on the removal of breathing motion and cardiac pulsatility connectome profiles.

Finally, we evaluated the addition of model-based nuisance regressors to the preprocessing strategies. Specifically, we added the physiological regressors used to model SLFOs (Kassinopoulos and Mitsis, 2019b), cardiac pulsatility and breathing motion (Glover et al., 2000). We found that including the regressor that models SLFOs reduces their effect on static FC for all preprocessing strategies apart from WM_50_ and WM_200_, but, in contrast to GSR, the similarity remains well above chance levels (**Supp. Fig. 3A**). Including the RETROICOR regressors related to breathing motion considerably reduced the breathing motion signature in the raw data; however, none of the preprocessing strategies benefited from including these regressors (**Supp. Fig. 3C**). On the contrary, including the RETROICOR regressors related to cardiac pulsatility completely removed the effect of the latter for the raw data and the MildA and MildB strategies (**Supp. Fig. 3D**), suggesting that conservative preprocessing strategies greatly benefit by adding the model-based regressors for cardiac pulsatility.

### 2.4 Connectome-based identification of individuals

We next investigated the extent to which FC matrices associated to physiological processes and head motion can identify an individual subject, and whether the accuracy of connectome-based fingerprinting is inflated by the examined nuisance processes (see Materials and methods – Connectome-based identification of individual subjects).

We initially considered all the edges of the FC matrices for subject identification (Gordon atlas: 40,755 edges). Accuracy was above chance for all database-target combinations for the nuisance processes, with rates up to 40% (**Figure 3A**). Breathing motion exhibited an intriguing bimodal distribution: database-target pairs that had the same phase encoding yielded much higher identification rates than database-target pairs with different phase encodings, even if the latter were acquired on the same day. This effect was also observed, although to a lesser extent, for cardiac pulsatility and head motion.

**Figure 3.**
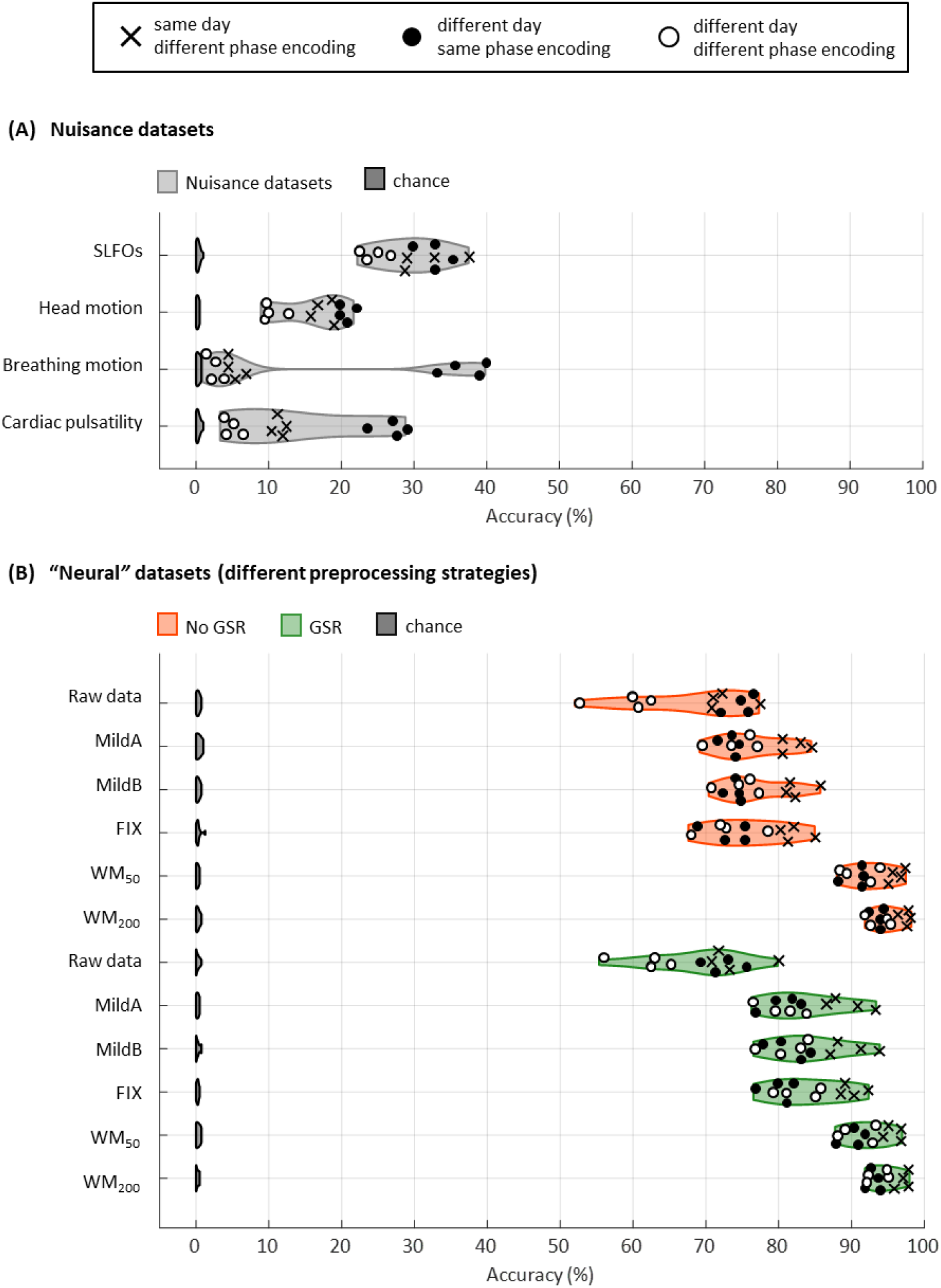
Connectome fingerprinting results. (**A**) Fingerprinting accuracy obtained using the static FC matrices from the generated nuisance datasets whereby non-neural fluctuations were isolated from the BOLD data. Above-chance level accuracy values were obtained for all nuisance processes, suggesting some degree of subject specificity in whole-brain connectivity profiles arises from nuisance fluctuations. (**B**) Fingerprinting accuracy obtained using the static FC matrices generated from each of the preprocessing strategies evaluated. The pairs of resting-state scans are indicated with different symbols, depending on whether they belong to the same or different day session, as well as whether they have the same phase encoding. Higher fingerprinting accuracy values were observed for white matter denoising approaches (WM_50_, WM_200_) compared to milder pipelines and FIX denoising. Both mild and more aggressive pipelines yielded higher subject discriminability for pairs of scans acquired on the same day. GSR increased the fingerprinting accuracy of milder strategies and FIX denoising.

Identification accuracy was much higher for the “neural” datasets compared to the nuisance datasets, with rates ranging from 52% to 99% (**Figure 3B**). The MildA, MildB and FIX techniques considerably improved the accuracy compared to the raw data, and the WM_50_ and WM_200_ techniques significantly outperformed all other preprocessing strategies. GSR considerably improved identification accuracy for the MildA, MildB and FIX strategies. Furthermore, we observed that for the raw data, database-target pairs from different days with the same phase encoding showed identification rates as high as the ones from the same day but different phase encoding. In contrast, for all other preprocessing strategies the database-target pairs from the same day were always higher.

We subsequently tested identification accuracy on the basis of within and between edges of specific functional networks to examine whether certain functional connections had a more pronounced contribution to individual subject discriminability. Results for the nuisance datasets are shown in **Figure 4A**, where it can be seen that nuisance processes yielded a markedly lower identification accuracy when using specific edges compared to using all edges (**Figure 3A**). Furthermore, functional connections between networks seemed to contribute more to the subject discriminability of SLFOs compared to connections within brain networks (*p*<0.001, Wilcoxon rank-sum).

**Figure 4.**
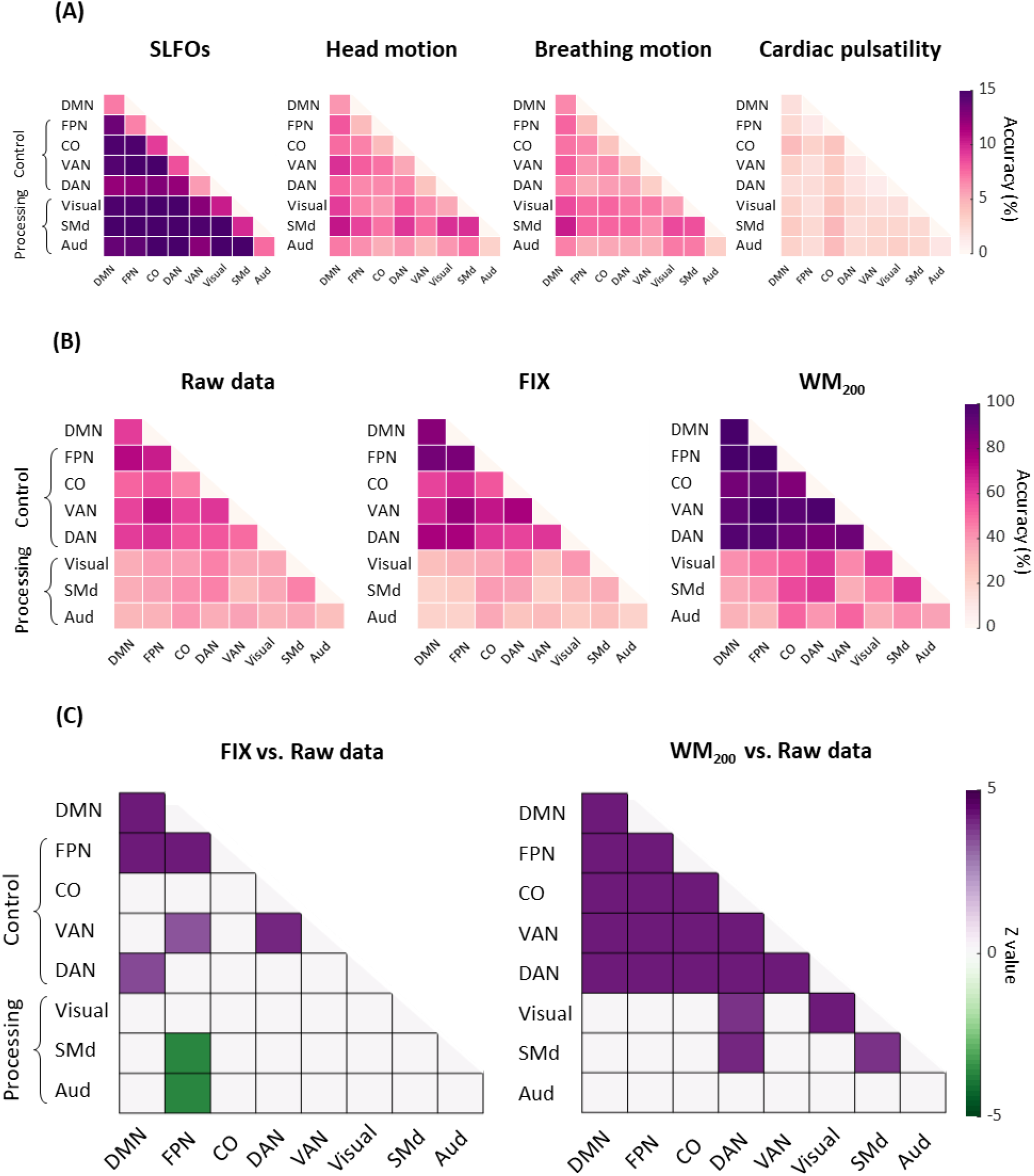
Connectome fingerprinting results using edges within and between networks. (**A**) Fingerprinting accuracy for SLFOs, head motion, breathing motion and cardiac pulsatility, averaged across all database-target pairs. (**B**) Fingerprinting accuracy for the raw data, FIX and WM_200_ pipelines, averaged across all database-target pairs. (**C**) Significant differences in fingerprinting accuracy obtained when using the FIX and WM_200_ pipelines as compared to the raw data (*p*<0.05, Bonferroni corrected, Wilcoxon rank-sum). Connectivity profiles within and between top-down control networks (FPN, CO, VAN, DAN) and DMN yielded higher identification accuracy compared to connectivity profiles within and between sensorimotor processing networks (Visual, SMd, Aud).

Regarding the “neural” datasets, we focused on the most aggressive strategies, namely FIX and WM_200_. Networks of “top-down” control (FPN, CO, DAN, VAN), as well as the DMN, yielded higher identification accuracy compared to sensorimotor processing networks (Visual, SMd, Aud) for all preprocessing strategies (**Figure 4B**). These results indicate that FC patterns in higher-order association cortices (“top-down” control networks) tend to be distinctive for each individual, whereas primary sensory and motor regions (processing networks) tend to exhibit similar patterns across individuals, consistent with previous studies (Finn et al., 2015; Gratton et al., 2018; Horien et al., 2019). We then tested for differences in identification accuracy between the raw data vs. FIX and WM_200_ data (**Figure 4C**, *p*<0.05, Bonferroni corrected, Wilcoxon rank-sum). FIX denoising significantly increased the subject discriminability of connections within and between several top-down control networks, but significantly decreased the subject discriminability of connections between the FPN and SMd, as well as the FPN and Aud networks. Conversely, WM_200_ denoising significantly increased the subject discriminability of connections within and between all top-down control networks, connections within the Visual and SMd networks, and connections of the DAN with the Visual and SMd network.

### 2.5 Physiological and head motion signatures in time-varying FC and the effect of preprocessing strategies

To examine the effect of physiological processes and head motion on time-varying FC estimates, we computed functional connectivity dynamics (FCD) matrices (Hansen et al., 2015) using the generated nuisance and “neural” datasets from each scan, whereby each FCD matrix captures the temporal evolution of FC patterns within a scan. We subsequently computed the similarity of the nuisance and “neural” FCD matrices at the individual level to examine the capability of various preprocessing strategies to reduce the confounds introduced by physiological processes and head motion on time-varying FC. A distribution of the similarity values, in this case Pearson’s correlation coefficients, is shown in **Figure 5** for each preprocessing strategy and nuisance process. We observed that the temporal evolution of FC patterns from SLFOs and head motion were similar to the ones observed in the raw data. An illustration of this similarity is shown in **Figure 6** for six subjects. On the other hand, breathing motion and cardiac pulsatility FCD matrices did not show similarities with the FCD matrices obtained from “neural” datasets (**Figure 5**).

**Figure 5.**
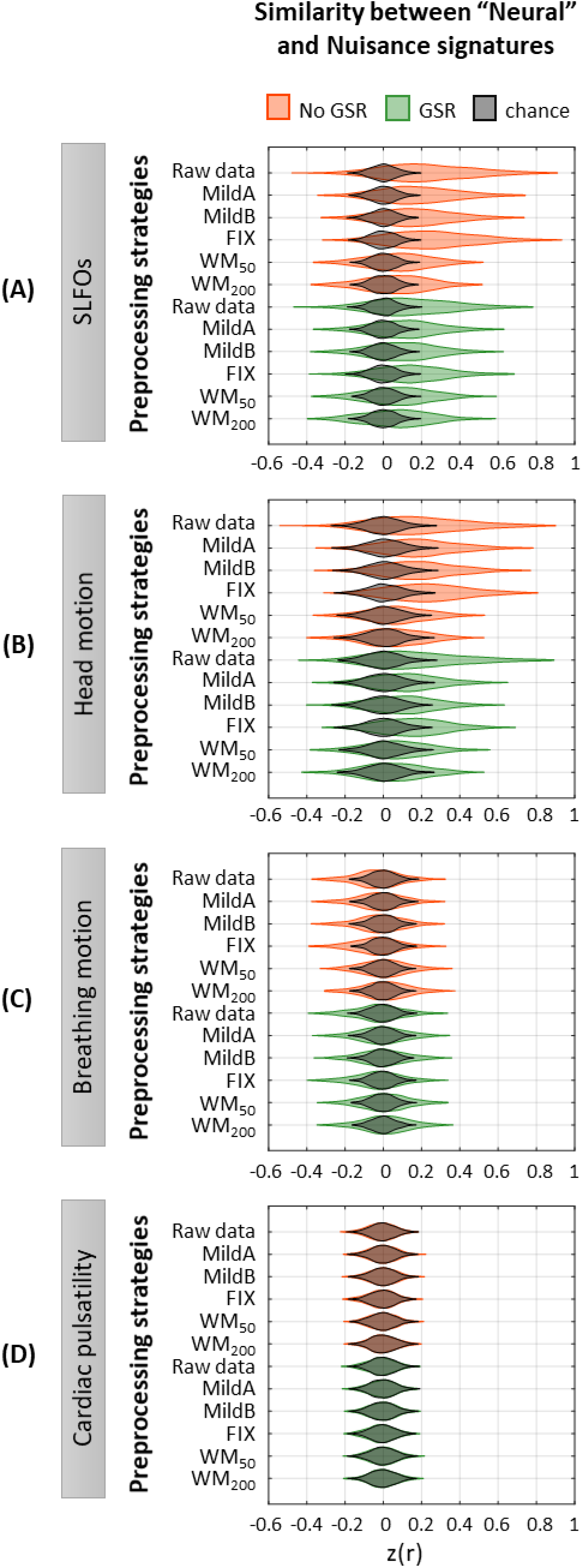
Effectiveness of preprocessing strategies in reducing functional connectivity dynamics (FCD) profiles induced by physiological and motion processes. Distribution of Pearson correlation coefficients across all 1,568 scans between the “neural” functional connectivity dynamics (FCD) matrix after each preprocessing pipeline and nuisance FCD matrices associated to (**A**) SLFOs, (**B**) head motion, (**C**) breathing motion, and (**D**) cardiac pulsatility. Correlation values were Fisher z transformed. Results shown in the top row of each subpanel (raw data) suggest that SLFOs and head motion most severely confound the FC matrices, whereas breathing motion and cardiac pulsatility do not induce artifactual dynamics. None of the examined strategies completely eliminated these effects.

**Figure 6.**
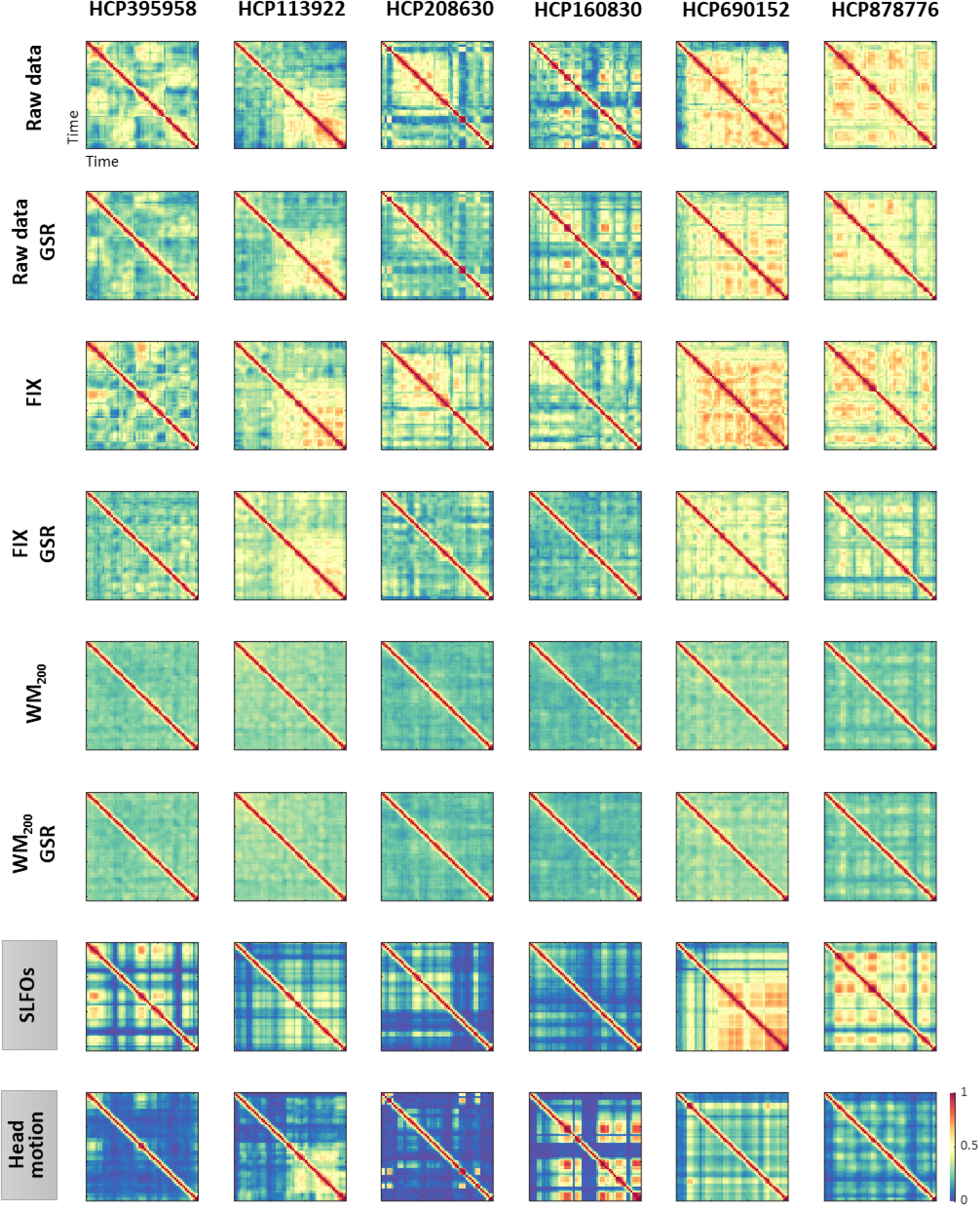
Functional connectivity dynamics (FCD) profiles associated with SLFOs and head motion resemble patterns commonly attributed to neural processes. Illustrative examples of FCD matrices from specific HCP subjects as obtained from the fMRI data for several pre-processing pipelines (rows 1-6), as well as from SLFOs and head motion (rows 7 and 8, respectively). All the examples are from the HCP scan Rest1_LR. These examples show a clear resemblance between FCD matrices computed from the “neural” datasets and the nuisance processes (SLFOs and head motion).

None of the preprocessing pipelines was able to vanish the effects of SLFOs and head motion. However, these effects were considerably reduced by the WM_50_ and WM_200_ strategies (**Figure 5A-B, Figure 6**). FIX denoising was the least successful strategy in terms of reducing the SLFOs’ signature (**Figure 5A, Figure 6**), similarly to static FC (**Figure 2E**), and only achieved the same levels of performance as other strategies after GSR. However, even after GSR none of the strategies reached chance levels (**Figure 5A**), in contrast with the static FC results (**Figure 2E**). GSR led also to a slight reduction in the similarity between the head motion and “neural” FCD matrices (**Figure 5B**), as in the case of static FC (**Figure 2F**).

## 3 Discussion

In this work, we characterized the effects of physiological processes and head motion on static and time-varying estimates of functional connectivity measured with BOLD fMRI. While the BOLD signal is considered a proxy of neural activity via changes in local blood oxygenation, physiological processes and motion artifacts can also induce variations in the BOLD signal, which can in turn lead to confounds in estimates of functional connectivity. Here, we developed an innovative framework to characterize the spatial signature of head motion and physiological processes (cardiac and breathing activity) on estimates of functional connectivity. Our results demonstrated that functional connectivity measures can be influenced by non-neural processes. Specifically, we identified stereotyped whole-brain functional connectivity profiles for SLFOs, head motion and breathing motion (**Figure 2A-C**), suggesting that these processes introduce a systematic bias in estimates of functional connectivity if they are not properly accounted for. Furthermore, we provided evidence that recurring patterns in time-varying FC can be attributed, to some extent, to SLFOs and head motion (**Figure 5, Figure 6**). We also assessed the performance of several state-of-the-art preprocessing strategies in mitigating the effects of nuisance processes, and showed that more aggressive preprocessing strategies such as FIX (Salimi-Khorshidi et al., 2014) and WM denoising (Kassinopoulos and Mitsis, 2019a) combined with GSR were the most effective with regards to removing the effects of non-neural processes for both static and time-varying FC analyses (**Figure 2E-H, Figure 5, Figure 6**). Finally, we evaluated the potential subject specificity of the connectivity profiles associated with physiological and motion confounds, along with their role as hypothetical contributors to connectome fingerprinting accuracy. Interestingly, we found that these non-neural functional connectivity patterns are to some extent subject specific (**Figure 3A**); however, fMRI data corrected for these confounds increased identification accuracy in connectome fingerprinting (**Figure 3B**), suggesting that the inter-individual differences in FC that facilitate subject identification are strongly neural and do not largely stem from physiological processes or head motion.

### 3.1 Spatially heterogeneous contributions of nuisance processes to the BOLD signal

It is well established that head and breathing motion affect areas at the edges of the brain (Jo et al., 2010; Patriat et al., 2015; Satterthwaite et al., 2013), whereas cardiac pulsatility affects areas near the large cerebral arteries just above the neck (Glover et al., 2000; Kassinopoulos and Mitsis, 2020). These observations are based on studies that typically examine the brain regions affected by the aforementioned sources of noise on a voxel-wise basis. However, at the voxel level we cannot easily assess whether the average fMRI signal from atlas-based ROIs includes significant contributions from these nuisance processes. In principle, it could be the case that the dynamics of artifacts associated with a specific nuisance process demonstrate significant variability across voxels, and as a result their effects cancel out when averaging voxels within an ROI. In the present study, we assessed the impact and regional variation of these nuisance processes in the Gordon parcellation (Gordon et al., 2016), a widely used atlas in the literature.

SLFOs related to changes in heart rate and breathing patterns were found to affect mostly sensory regions including the visual and somatosensory cortices (particularly of the face) (**Figure 1A**), which correspond to regions with a high density of veins (Bernier et al., 2018; Huck et al., 2019). The spatial pattern of SLFOs is very similar to statistical maps reported in prior works, which have highlighted brain regions highly correlated with the global signal (Billings and Keilholz, 2018; Glasser et al., 2016; Li et al., 2019a; Power et al., 2017; Tong et al., 2013; Zhang et al., 2019). This is not surprising, since the global signal is strongly driven by fluctuations in heart rate and breathing patterns (Birn et al., 2006; Chang and Glover, 2009a; Falahpour et al., 2013; Kassinopoulos and Mitsis, 2019b; Shmueli et al., 2007). Moreover, we show evidence that SLFOs and cardiac pulsatility do not affect the same brain regions, consistent with (Chen et al., 2019; Kassinopoulos and Mitsis, 2019b; Tong and Frederick, 2014). Specifically, cardiac pulsatility was more dominant in regions such as the insular and auditory cortices, which align with cortical branches of the middle cerebral artery (**Figure 1D**) and are the regions with highest arterial density (Bernier et al., 2018).

Regarding head motion, previous studies found that its effect was more pronounced in prefrontal, sensorimotor and visual brain regions (Satterthwaite et al., 2013; Yan et al., 2013). However, these studies did not remove breathing artifacts from the realignment parameters, which are present even in single-band datasets (Gratton et al., 2020), and thus were unable to disentangle whether a specific type of motion affected particular brain regions. In the present work, we regressed out breathing motion from the realignment parameters, and observed that sensorimotor and visual areas were strongly affected by head motion (**Figure 1B**), whereas breathing motion artifacts were more pronounced in the prefrontal cortex and brain regions in the parietal and temporal cortices (**Figure 1C**). Furthermore, Yan et al. showed that framewise displacement was positively correlated with sensory regions and negatively correlated with prefrontal regions. Collectively, these findings suggest that most regions exhibit an increase in the BOLD signal due to head and breathing motion, whereas the prefrontal cortex may exhibit a decrease in the BOLD signal likely due to breathing-related chest movements.

### 3.2 Physiological and head motion signatures in static FC

Head motion is considered the biggest source of confound for FC fMRI studies and there is a significant effort from the neuroimaging community towards developing and evaluating preprocessing strategies that mitigate its effects (Ciric et al., 2017; Parkes et al., 2018; Power et al., 2015). On the other hand, while it has been shown that SLFOs affect the default-mode network (Birn et al., 2014, 2008a; Chang and Glover, 2009a), and high frequency cardiac and breathing artifacts influence the BOLD signal (Glover et al., 2000; Power et al., 2019), a systematic investigation of the effects of physiological processes in the context of whole-brain FC is lacking in the literature. In the present study, we evaluated collectively the impact of the aforementioned sources of noise on whole-brain fMRI resting-state FC.

Our results revealed that all four nuisance datasets exhibited mainly positive correlations between ROIs (**Figure 2A-D**), suggesting that the presence of nuisance fluctuations in a conventional fMRI dataset typically leads to a shift of correlation values towards more positive numbers. In other words, in the case of an fMRI dataset that has not been corrected for nuisance fluctuations, two ROIs for which neural-related fluctuations are negatively correlated could be found to be positively correlated due to the presence of similar nuisance fluctuations in the ROIs. Furthermore, we observed that SLFOs and head motion confounded FC to a larger degree compared to breathing motion and cardiac pulsatility (**Figure 2E-H**).

Our results suggest that SLFOs due to spontaneous changes in heart rate or breathing patterns inflate connectivity (towards more positive values) across the whole brain but particularly for edges within the visual network, as well as edges between the visual and the rest of the networks (**Figure 2A**). It is well known that the visual cortex is characterized by the highest venous density (Bernier et al., 2018), possibly due to its functional importance (Collins et al., 2010). In addition, it has been shown that brain regions with higher vascular density exhibit larger amplitude of spontaneous BOLD fluctuations (Vigneau-Roy et al., 2014). Therefore, it is likely that the structure of the SLFOs’ connectome profile may largely reflect the underlying vascular architecture. The effect of SLFOs on static FC was considerably reduced after WM denoising, while additionally performing GSR almost removed this effect (**Fig. 4E**). Notably, FIX denoising without GSR was unable to remove the confounds introduced by SLFOs, which is consistent with recent studies showing that global artifactual fluctuations are still prominent after FIX denoising (Burgess et al., 2016; Glasser et al., 2018; Kassinopoulos and Mitsis, 2019b; Power et al., 2018, 2017).

Head motion was found to influence the connectivity within the visual and sensorimotor networks (**Figure 2B**), in line with previous studies (Power et al., 2012; Satterthwaite et al., 2012; van Dijk et al., 2012). Our results showed that only regressing out the realignment parameters and average WM/CSF signals (with or without expansion terms) is not sufficient to remove the effects of head motion (**Figure 2F**, MildA and MildB pipelines), which is consistent with findings in (Parkes et al., 2018). Among all preprocessing strategies, WM denoising yielded the largest reduction of motion effects (**Figure 2F**). The two pipelines WM_50_ and WM_200_ refer to the removal of 50 and 200 white matter regressors from the data (i.e. principal components obtained from the white matter compartment). In our previous study we showed that while both pipelines yielded high large-scale network identifiability compared to other pipelines, the more aggressive WM_200_ resulted in a larger reduction of motion artifacts compared to WM_50_ (Kassinopoulos and Mitsis, 2019a). The results of the current study also show a stronger reduction of head motion effects for the former compared to the latter (**Figure 2F**), which may explain the higher accuracy in connectome fingerprinting observed for the pipeline WM_200_ compared to WM_50_ (**Figure 3B**).

A natural concern regarding the head motion connectome profile is that it may reflect motor-related activity (Yan et al., 2013). Even though motor-related neural activity would be expected to lag the instantaneous motion traces due to the sluggishness of the hemodynamic response, we cannot exclude the scenario that the head motion connectome profile reflects the neural correlates of the executed movements and eye adjustments to fixate on the cross. Nonetheless, even if preprocessing strategies remove neural activity associated with spontaneous head movements, this source of neural activity is typically of no interest in resting-state fMRI studies.

Furthermore, we provide evidence that head and breathing motion do not affect functional connectivity in the same manner. Specifically, breathing motion was found to inflate within-hemisphere connectivity (**Figure 2C**). This bias seems to arise as a result of factitious motion rather than real motion of the head, since it is related to the LR/RL phase encoding direction (see section 4.4 for more details). All preprocessing strategies yielded a substantial reduction of artifacts related to breathing motion, with FIX denoising being the most effective (**Figure 2G**).

In our dataset, cardiac pulsatility did not seem to have a large effect on FC, neither in cortical nor subcortical regions (**Figure 2D, Supp. Fig. 5D**), and its effect was entirely removed with more aggressive pipelines such as FIX and WM denoising (**Figure 2H**), as well as with model-based techniques (**Supp. Fig. 3D**). However, it has been recently reported that the 3T HCP dataset has poor temporal signal-to-noise ratio in the subcortex (Ji et al., 2018; Seitzman et al., 2019b). Therefore, it is possible that we may have underestimated the effect of cardiac pulsatility in functional connections involving subcortical regions.

It is important to note that our proposed methodology assumes that the stereotyped nuisance connectome profiles do not resemble the true neural connectome profiles. However, in principle, nuisance fluctuations could give rise to similar spatial patterns as neurally-driven fluctuations. A recent study by (Bright et al., 2018) provided evidence that physiological fluctuations (end-tidal CO_2_) give rise to networks that spatially resemble neurally-driven networks linked to working memory and visual stimuli. The authors suggested that this phenomenon may be due to the vasculature adapting to the neural network architecture, as vascular and neuronal growth processes evolve concurrently during development (Quaegebeur et al., 2011). These findings suggest a possible caveat of our methodology when assessing pre-processing strategies, as pipelines that yield the lowest similarity between nuisance and “neural” FC matrices might also remove some signal of interest. Nonetheless, the pre-processing strategies that were found in this study to reduce the nuisance effects the most (i.e. FIX and WM denoising combined with GSR) have been shown to demonstrate the highest improvement in large-scale network identifiability in an earlier study (Kassinopoulos and Mitsis, 2019a). In addition, these pipelines were found to exhibit the highest accuracy in connectome fingerprinting (**Figure 3B**). These results suggest that they are able to adequately remove the effects of nuisance processes while also preserving the signal of interest.

### 3.3 Physiological and head motion signatures in time-varying FC

The investigation of neural dynamics using resting-state fMRI is a promising avenue of research that has gained increasing attention lately (Hutchison et al., 2013; Lurie et al., 2019). Yet, there is skepticism regarding its validity and underlying origins. For instance, variations in FC over shorter time-scales (i.e. minutes) could largely be explained by sampling error, acquisition artifacts and subject arousal (Hindriks et al., 2016; Laumann et al., 2017; Savva et al., 2019), as well as head motion and physiological processes (Nalci et al., 2019; Nikolaou et al., 2016).

In the present study, we sought to evaluate whether non-neural fluctuations could partly explain the recurrent connectivity patterns observed in fMRI studies. To this end, we computed time-resolved FC dynamics (Hansen et al., 2015) for all four nuisance datasets, and assessed their similarity with time-resolved FC dynamics obtained from fMRI data preprocessed employing widely used denoising strategies (“neural” datasets). The FC dynamics of head motion and SLFOs datasets were markedly similar to the FC dynamics observed in preprocessed fMRI data (**Figure 5A-B**), albeit the similarity was smaller compared to static FC (**Figure 2E-F**). When observing the time-resolved FC matrices (**Figure 6**), it becomes apparent that a large component of variability in FC patterns is due to non-neural processes, and that these patterns remain after implementing popular preprocessing pipelines such as MildA, MildB and FIX. These results are aligned with the observation that even after regressing out nuisance processes from the BOLD signal, correlations between time-varying FC measures and nuisance fluctuations remain (Nalci et al., 2019). WM denoising was found to be the most efficient strategy in terms of mitigating the influence of nuisance processes on time-varying FC (**Figure 5A-B**).

After WM denoising, variability in FC patterns was greatly diminished (**Figure 6**), even when those patterns could not be directly associated with any nuisance process (**Supp. Fig. 4**). These results can be interpreted in two ways that are not mutually exclusive: (1) A significant fraction of the variability in FC patterns is a result of non-neural confounds and WM denoising is able to remove most of these confounds. This is supported by the fact that many other nuisance processes, which we did not examine here (e.g. arterial blood pressure, CO_2_ concentration, scanner instabilities), can influence the BOLD signal and time-varying FC patterns (Nikolaou et al., 2016; Whittaker et al., 2019; Wise et al., 2004). (2) WM denoising removes a considerable fraction of variance of neural origin. Future work with concurrent direct measurements of neuronal activity (e.g. electroencephalography, calcium imaging) and additional physiological recordings would be instrumental for resolving to which extent time-varying FC is the result of underlying neural dynamics.

While head motion and SLFOs were found to be strongly associated to recurrent connectivity patterns, breathing motion and cardiac pulsatility do not seem to be a main concern for time-varying FC studies (**Figure 5C-D**). Likely, the effects of breathing motion and cardiac pulsatility do not influence time-varying FC, because their effect on the BOLD signal does not change from window to window, possibly due to their quasi-periodic nature. In contrast, the levels of head motion vary across time windows, which can modulate time-varying FC patterns. Heart rate and breathing patterns can be relatively constant during some time periods, whereby SLFOs are not expected to influence the BOLD signal and, in turn, the FC measures across time windows. On the other hand, in other instances heart rate and breathing patterns may change considerably over time, whereby SLFOs are expected to influence the BOLD signal and thus modulate the FC measures across time windows. In other words, ROIs sensitive to head motion and SLFOs are likely to exhibit a time-varying signal-to-noise ratio depending on the presence of these sources of noise, which eventually leads to confounds in time-varying FC measures.

Importantly, none of the evaluated pipelines were able to completely remove these confounds. It was only recently that researchers have started to examine the performance of pre-processing pipelines in the context of time-varying FC (Lydon-Staley et al., 2019), albeit with a focus on motion effects, thus more work is needed to identify effective data cleaning strategies for resting-state time-varying FC studies.

### 3.4 Global signal regression

The practice of removing the GS from fMRI data (i.e. GSR) has been adopted by many fMRI investigators as it has been linked to head motion artifacts and fluctuations in heart rate and breathing patterns (Birn et al., 2006; Byrge and Kennedy, 2018; Chang and Glover, 2009a; Falahpour et al., 2013; Kassinopoulos and Mitsis, 2019b; Power et al., 2018, 2014; Shmueli et al., 2007). Further, GSR has been shown to increase the neuronal-hemodynamic correspondence of FC measures extracted from BOLD signals and electrophysiological high gamma recordings (Keller et al., 2013), as well as strengthen the association between FC and behaviour (Li et al., 2019b). On the other hand, studies capitalizing on EEG-fMRI data have reported an association between the GS amplitude and vigilance measures (Wong et al., 2016, 2013) and individual differences in the global signal topography have been related to behavior and cognition (Li et al., 2019a). Thus, as there is evidence that GSR may remove neuronal-related activity in addition to nuisance-related fluctuations, GSR still remains a controversial pre-processing step (Liu et al., 2017; Murphy et al., 2009; Murphy and Fox, 2017).

Our results provide evidence that, in the context of static FC, GSR removes physiological fluctuations related to SLFOs and to a lesser extent head motion artifacts (**Figure 2E-F**). Note that GSR does not account for breathing motion artifacts (**Figure 2G**) but rather changes in breathing patterns and deep breaths, which are related to SLFOs and possibly the head motion component at ~0.12 Hz (Power et al., 2019). Furthermore, GSR improved connectome fingerprinting accuracy (**Figure 3B**), which suggests that by removing nuisance fluctuations due to SLFOs and head motion, GSR enhances the individual specificity of connectivity profiles. Overall, our results suggest that the strong reduction in the effects of SLFOs and head motion achieved by GSR outweighs the possible loss of neuronal-driven fluctuations when examining FC patterns. GSR is particularly important when using ICA-based noise correction techniques such as FIX and AROMA (Pruim et al., 2015; Salimi-Khorshidi et al., 2014), since ICA components related to SLFOs frequently exhibit similar spatial patterns and frequency profile to neural components and thus are classified as non-artifactual and remain in the data after denoising.

Regarding time-varying FC, GSR did not reduce the effect of nuisance processes equally well compared to static FC (**Figure 2E-F vs. Figure 5A-B**). Nonetheless, a recent study evaluating preprocessing strategies in the context of time-varying FC showed that incorporating GSR in the preprocessing improved the identification of modularity in functional networks (Lydon-Staley et al., 2019). This may indicate that GSR was able to remove nuisance processes that we did not evaluate in the current study. These processes may be related to scanner instabilities, CO_2_ concentration (Power et al., 2017; Wise et al., 2004) and finger skin vascular tone (Kassinopoulos and Mitsis, 2020; Özbay et al., 2019), which are known to be reflected on the GS.

Despite the effectiveness of GSR in reducing nuisance confounds from the data, we cannot exclude the possibility of removing some neuronal-related fluctuations. Alternatives to GSR that have been proposed to remove global artifacts include time delay analysis using “rapidtide” (Tong et al., 2019), removal of the first principal component from the fMRI data (Carbonell et al., 2011), removal of fluctuations associated to large clusters of coherent voxels (Aquino et al., 2020), and the use of temporal ICA (Glasser et al., 2018), albeit the latter is only applicable to datasets with a large number of subjects such as the HCP.

### 3.5 The effect of phase encoding direction in connectivity

Earlier studies have demonstrated that chest wall movements due to breathing perturb the B0 field (Raj et al., 2001, 2000; Van de Moortele et al., 2002), which has consequences on EPI fMRI data. While this phenomenon is not fully understood, it seems to have two main effects that are observable along the phase encoding direction: (1) Breathing causes factitious motion of the fMRI volumes in the phase encoding direction (Raj et al., 2001, 2000). This effect has sparkled attention recently, since it has been recognised that it may have critical implications for motion correction when performing censoring (i.e. removal of motion-contaminated fMRI volumes) in multi-band (Fair et al., 2019; Power et al., 2019) and single-band (Gratton et al., 2020) data. (2) Breathing induces artifacts on voxel timeseries that depend on the location of those voxels along the phase encoding direction (Raj et al., 2001, 2000). Our results provide further evidence in support of the latter effect. Specifically, we found that depending on the phase encoding direction (LR or RL), breathing motion artifacts were more pronounced in the left or right hemisphere respectively (**Figure 1C**). Moreover, we observed that breathing motion increased within-hemisphere connectivity for both phase encoding scan types (**Figure 2C, Supp. Fig. 1C**), which implies that breathing induces artifactual fluctuations that are to a certain extent different between hemispheres. However, note that the connectome profile of breathing motion exhibited some differences between the two phase encoding directions (**Supp. Fig. 1C**), which explains the higher connectome fingerprinting accuracy in the breathing motion dataset when examining pairs of scans with the same phase encoding direction, compared to scans with different phase encoding direction (**Figure 3A**).

Our results point to a systematic effect of breathing on static FC through variations in the B0 magnetic field. Importantly, this systematic bias is contingent on the phase encoding direction, which seems to indicate that factitious rather than real motion is the predominant source of respiration-related motion artifacts in fMRI, as has been previously suggested (Brosch et al., 2002; Raj et al., 2001). Even though common preprocessing pipelines greatly reduce these effects, they do not eliminate them (**Figure 2G**). Thus, studies that consider datasets with different phase encodings, should be aware of the effect of phase encoding on FC, especially if data from different groups have been acquired with different phase encodings.

### 3.6 Individual discriminability

Test-retest reliability is important for establishing the stability of inter-individual variation in fMRI FC across time. However, apart from neural processes, nuisance processes can also have an impact on test-retest reliability, given the subject-specific nature of physiological processes (Batchvarov et al., 2002; Golestani et al., 2015; Malik et al., 2008; Pinna et al., 2007; Pitzalis et al., 1996; Power et al., 2020; Reland et al., 2005) and head motion (van Dijk et al., 2012; Zeng et al., 2014). This leads to the concerning notion that nuisance processes may be artifactually driving the reports of high reliability in FC measures. For instance, it has been reported that the median of intraclass correlation values across functional connections, which is a metric of test-retest reliability, is reduced when a relatively aggressive pipeline is used (Birn et al., 2014; Parkes et al., 2018). Furthermore, motion can classify subjects at above-chance levels (Horien et al., 2019), and breathing motion is more prominent in older individuals and those with a higher body mass index (Gratton et al., 2020). In the present study, we examined the potential effect of nuisance processes on subject discriminability using connectome fingerprinting.

#### 3.6.1 Whole-brain identification

To assess the individual discriminability of nuisance processes, we performed connectome fingerprinting analysis using the generated nuisance datasets. All nuisance processes exhibited identification accuracy above chance level (**Figure 3A**). Pairs of scans with the same phase encoding yielded higher identification accuracy than pairs of scans with different phase encoding (**Figure 3A**). This effect is particularly evident for breathing motion, and to a lesser extent cardiac pulsatility and head motion. This observation suggests that not only these confounds exert a distinctive artifactual spatial pattern that is dependent on the phase encoding direction, which can be also observed upon careful examination of **Figure 1B-D**, but also that this artifactual pattern is to a certain degree subject-specific. On the other hand, the subject discriminability of SLFOs is not modulated by phase encoding (**Figure 3A**). Given the nature of SLFOs (i.e. they affect the BOLD signal through changes in CBF), the high subject discriminability of SLFOs suggests a certain degree of idiosyncrasy that is possibly related to the vascular architecture of an individual. Overall, our results suggest that there is some degree of subject discriminability in nuisance processes.

Identification accuracies of “neural” datasets were very high for all preprocessing strategies (**Figure 3B**), in line with previous studies (Finn et al., 2015; Horien et al., 2019). WM denoising, which was found to be the most effective strategy for reducing confounds due to head motion and physiological fluctuations (**Figure 2E-H**), yielded also the highest accuracy in connectome fingerprinting (**Figure 3B**), suggesting that the high subject discriminability observed in the HCP data is not due to the presence of confounds. Interestingly, the increased accuracy observed in the nuisance datasets for scans with the same phase encoding (**Figure 3A**) was also observed in the case of the raw data (**Figure 3B**). In contrast, for the rest of the pipelines the difference in accuracy between pairs of scans from different days with the same or different phase encoding direction vanishes (**Figure 3B**). This is likely because of the reduction of nuisance effects, mainly breathing motion artifacts. Note also that for both mild and aggressive pipelines, pairs of scans from the same day exhibited higher accuracies compared to pairs of scans from different days, which cannot be attributed to potential residuals of nuisance fluctuations (**Figure 3A**). Possible explanations for this finding are that the functional connectome of a subject reflects some aspects of their vigilance levels (Tagliazucchi and Laufs, 2014; Thompson et al., 2013a; Wang et al., 2016), mind-wandering (Gonzalez-Castillo et al., 2019; Gorgolewski et al., 2014; Kucyi, 2018; Kucyi and Davis, 2014), or the effect of time of day (Hodkinson et al., 2014; Jiang et al., 2016; Orban et al., 2020; Shannon et al., 2013), which can differ across sessions. Overall, the high connectome-based identification accuracies reported in the literature do not appear to be driven by nuisance confounds, suggesting a neural origin underpinning the inter-individual differences in connectivity. Nonetheless, it is worth pointing out that subject variability in the magnitude of functional connections has been shown to arise as a result of spatial topographical variability in the location of functional regions across individuals (Bijsterbosch et al., 2018), which could also explain the high subject discriminability observed in fMRI-based connectomes.

#### 3.6.2 Network-based identification

We observed that edges within association cortices (e.g. parts of the frontoparietal, default mode, and cinguloopercular systems) exhibited the highest subject specificity (**Figure 4B**), consistent with previous studies (Finn et al., 2015; Gratton et al., 2018; Horien et al., 2019; Mueller et al., 2013; Seitzman et al., 2019a; Vanderwal et al., 2017). The fact that association cortices are the most evolutionarily recent (Zilles et al., 1988) and are thought to be involved in higher-level functions (Cole et al., 2014, 2013; Dosenbach et al., 2007; Gratton et al., 2017; Raichle, 2015) has been posited as a possible reason for the high identification accuracy yielded by these networks. On the other hand, it has also been speculated that medial frontal and frontoparietal networks exhibit the highest identification accuracy as a result of being less prone to distortions from susceptibility artifacts (Horien et al., 2019; Noble et al., 2017). If the latter was the case, we would expect to see decreased accuracy for these networks when probing the nuisance datasets. However, we did not observe such a tendency for any of the nuisance processes evaluated (**Figure 4A**), and preprocessing strategies that successfully removed nuisance processes yielded enhanced subject discriminability of control networks and the DMN (**Figure 4C**). These results seem to indicate that the basis of the high identification rates for association cortices is of neural origin, and thus that resting-state fMRI-based connectome fingerprinting can capture idiosyncratic aspects of cognition reflected on the resting-state functional characteristics of the association cortex.

## 4 Conclusions

The current study introduces a novel framework for assessing the effects of the main fMRI confounds on static and time-varying FC. Our results suggest that head motion and systemic BOLD fluctuations associated to changes in heart rate and breathing patterns cause systematic biases in static FC and result in recurrent patterns in time-varying FC. Data-driven techniques based on decomposing the data into principal or independent components (PCA, ICA), combined with GSR, lead to the strongest reduction of the aforementioned effects. Importantly, these preprocessing strategies also improve connectome-based subject identification, indicating that the high subject discriminability reported in the literature is not attributable to nuisance processes.

## 5 Materials and methods

### 5.1 Human Connectome Project (HCP) dataset

The resting-state fMRI data analysed in this study are from the S1200 release of the 3T HCP dataset (Smith et al., 2013; Van Essen et al., 2013), which consists of young, healthy twins and siblings (age range: 22-36 years). The HCP dataset includes, among others, resting-state data acquired on two different days, during which subjects were instructed to keep their eyes open and fixated on a cross-hair. Each day included two consecutive 15-min restingstate runs, acquired with left-to-right (LR) and right-to-left (RL) phase encoding direction. During each fMRI run, 1200 frames were acquired using a gradient-echo echo-planar imaging (EPI) sequence with a multiband factor of 8, spatial resolution of 2 mm isotropic voxels, and a TR of 0.72 s. Further details of the data acquisition parameters can be found in previous publications (Smith et al., 2013; Van Essen et al., 2012). Concurrently with fMRI images, cardiac and respiratory signals were measured using a standard Siemens pulse oximeter placed on the fingertip and a breathing belt placed around the chest, with a 400 Hz sampling rate.

We only considered subjects who had available data from all 4 runs, and excluded subjects based on the quality of the physiological recordings (see section 2.2.1 below for details). Pulse oximeter and respiratory belt signals from ~1000 subjects were first visually inspected to determine their quality, since their traces are often not of sufficient quality for reliable peak detection (Power, 2019). The selection process resulted in a final dataset with 392 subjects (ID numbers provided in Supp. Material).

### 5.2 Preprocessing

#### 5.2.1 Preprocessing of physiological recordings

After selecting subjects with good quality traces, the pulse wave was processed to automatically detect beat-to-beat intervals (RR), and the heart rate signal was further computed as the inverse of the time differences between pairs of adjacent peaks and converted to units of beats-per-minute (bpm). Heart rate traces were visually checked to ensure that outliers and abnormalities were not present. An outlier replacement filter was used to eliminate spurious changes in heart rate when these changes were found to be due to sporadic noisy cardiac measurements (for more details see **Supp. Figs. 1 and 2** from (Kassinopoulos and Mitsis, 2019b)). We also excluded subjects with a heart rate of exactly 48 bpm and lack of heartbeat interval variability, as they have been pointed out as outliers in recent studies (Orban et al., 2020; Valenza et al., 2019). The signal from the breathing belt was detrended linearly, visually inspected and corrected for outliers using a replacement filter. Subsequently, it was low-pass filtered at 5 Hz and *Z*-scored. The respiratory flow, proposed in (Kassinopoulos and Mitsis, 2019b) as a robust measure of the absolute flow of inhalation and exhalation of a subject at each time point, was subsequently extracted by applying further smoothing on the breathing signal (moving average filter of 1.5 sec window) and, subsequently, computing the square of the derivative of the smoothed breathing signal. Finally, heart rate and respiratory flow time-series were re-sampled at 10 Hz.

An example code (Preprocess_Phys.m) showing the detailed specifications of the algorithms used during the preprocessing of the physiological signals is available on github.com/mkassinopoulos/PRF_estimation/.

#### 5.2.2 Preprocessing of fMRI data: assessing the impact of nuisance correction strategies

From the HCP database we downloaded the minimally preprocessed data described in (Glasser et al., 2013) and the FIX-denoised data, both in volume and surface space. Briefly, the minimal preprocessing pipeline included removal of spatial distortion, motion correction via volume re-alignment, registration to the structural image, bias-field correction, 4D image intensity normalization by a global mean, brain masking, and non-linear registration to MNI space. Further steps to obtain surface data were volume to surface projection, multimodal inter-subject alignment of the cortical surface data (Robinson et al., 2014), and 2 mm (FWHM) surface-constrained smoothing. Additional steps following minimal preprocessing to obtain the FIX-denoised data were de-trending using a mild high-pass filter (2000 s), head motion correction via 24 parameter regression, and denoising via spatial ICA followed by an automated component classifier (FMRIB’s ICA-based X-noiseifier, FIX) (Griffanti et al., 2014; Salimi-Khorshidi et al., 2014). Minimal spatial smoothing (FWHM = 4 mm) was applied to the downloaded minimally preprocessed and FIX-denoised volumetric data. Both minimally preprocessed and FIX-denoised data were parcellated employing the Gordon atlas across 333 regions of interest (ROIs) (Gordon et al., 2016) and the Seitzman atlas across 300 ROIs (Seitzman et al., 2019b), using the surface and volume space data, respectively. ROIs that did not belong to a brain network were disregarded, hence a total of 286 ROIs (Gordon atlas) and 285 ROIs (Seitzman atlas) were retained for further analyses. The main differences between these two brain parcellations, apart from being computed on the surface and volume space respectively, are that the ROIs in the Gordon atlas do not have the same size, whereas in the Seitzman atlas the ROIs are all spheres of 8 mm diameter, and that the Gordon atlas only includes cortical regions, whereas the Seitzman atlas includes cortical and subcortical regions. The results from the Gordon atlas are presented in the main manuscript whereas the results from the Seitzman atlas can be found in the Supplementary Material (**Supp. Fig. 5, Supp. Fig. 6, Supp. Fig. 7**). Further, the parcellated data were high-pass filtered at 0.01 Hz.

In addition to the FIX-denoising strategy, several other data-driven preprocessing techniques were evaluated to assess the extent to which they were able to remove physiological and motion-driven confounds (**Table 1**). We chose pipelines that had been used in the landmark FC studies of (Finn et al., 2015) and (Laumann et al., 2017).

**Table 1.**
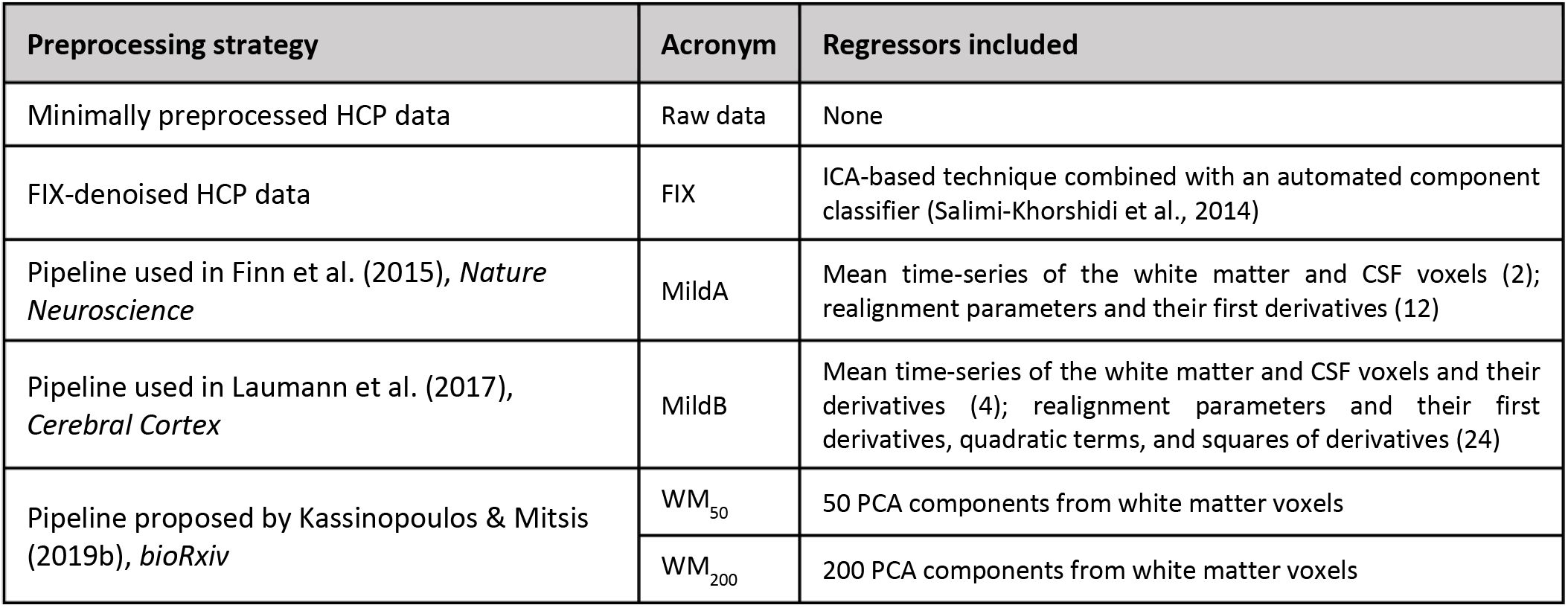
Preprocessing strategies examined. All strategies were evaluated with and without global signal regression (GSR).

These were denoted as “mild” pipelines, since they regress out considerably fewer components compared to FIX. Further, we also included two more aggressive pipelines that were found to outperform previously proposed techniques in terms of network identifiability (Kassinopoulos and Mitsis, 2019a). Nuisance regression was performed after the minimally preprocessed data had been parcellated to reduce computational time. All preprocessing strategies were evaluated with and without global signal regression (GSR), since the latter is still somewhat controversial (Liu et al., 2017; Murphy et al., 2009; Murphy and Fox, 2017). To facilitate the comparison between preprocessing strategies, the minimally preprocessed data were also evaluated, yielding in total 12 preprocessing strategies. Given that the minimal preprocessing pipeline consists of only the initial steps for fMRI denoising, for simplicity in the results we refer to these data as raw data. The regressors included in each preprocessing strategy can be found in Table 1. Note that for the pipeline from (Laumann et al., 2017) the derivative of the global signal was also regressed out. The global signal for the surface and volumetric data was computed as the average fMRI timeseries across vertices and the whole brain respectively.

### 5.3 Nuisance processes evaluated

The following four nuisance processes were considered (**Table 2**):

a. **Systemic low frequency oscillations (SLFOs):** SLFOs refer to non-neuronal global BOLD fluctuations. Major sources of SLFOs are spontaneous fluctuations in the rate or depth of breathing (Birn et al., 2006) and fluctuations in heart rate (Chang et al., 2009; Shmueli et al., 2007). The former mainly exert their effects via changing the concentration of arterial CO_2_, which is a potent vasodilator, altering CBF and thus the BOLD fMRI signal (Birn et al., 2008a, 2008b; Chang and Glover, 2009b; Prokopiou et al., 2019; Wise et al., 2004), Importantly, there is evidence that SLFOs are a more substantial source of physiological noise in BOLD fMRI compared to high frequency cardiac pulsatility and breathing motion artifacts (Tong et al., 2019; Tong and Frederick, 2014). In this study, SLFOs were modelled following a framework proposed in our previous work (Kassinopoulos and Mitsis, 2019a; scripts available on github.com/mkassinopoulos/PRF_estimation/). Briefly, the extracted heart rate and respiratory flow signals were fed into an algorithm that estimated scan-specific physiological response functions (PRFs). This algorithm estimates PRF curves that maximize the correlation between their convolution with heart rate and respiratory flow and the global signal of the same scan, while ensuring that the shapes of the PRF curves are physiologically plausible. The heart rate and respiratory flow signals were subsequently convolved with their respective PRFs and added to each other, yielding time-series that reflect the total effect of SLFOs (**Supp. Fig. 8**). These time-series were used in the current study as the physiological regressor related to SLFOs.
b. **Cardiac pulsatility:** Pulsatility of blood flow in the brain can cause pronounced modulations of the BOLD signal (Noll and Schneider, 1994), which tend to be localized along the vertebrobasilar arterial system and the sigmoid transverse and superior sagittal sinuses (Dagli et al., 1999; Kassinopoulos and Mitsis, 2019b). We modelled fluctuations induced by cardiac pulsatility using 6 regressors obtained with 3^rd^ order RETROICOR (Glover et al., 2000), based on the pulse oximeter signal of each scan.
c. **Breathing Motion:** Chest movements during the breathing cycle generate head motion in the form of head nodding by mechanical linkage through the neck, but also factitious motion (also known as pseudomotion) through small perturbations of the B0 magnetic field caused by changes in abdominal volume when air enters the lungs (Power et al., 2019; Raj et al., 2001; Van de Moortele et al., 2002). We modelled breathing-induced fluctuations using 6 regressors obtained with 3^rd^ order RETROICOR (Glover et al., 2000), based on the respiratory signal of each scan.
d. **Head motion:** Subject motion produces substantial signal disruptions in fMRI studies (Friston et al., 1996; Power et al., 2012) and is a major confound when evaluating connectivity differences between groups with dissimilar tendencies for motion (Makowski et al., 2019; Satterthwaite et al., 2012; van Dijk et al., 2012). We quantified head motion using the six realignment parameters as well as their temporal first derivatives provided by the HCP. The six physiological regressors related to breathing motion were regressed out from the realignment parameters and their derivatives, since true and factitious motion due to breathing is reflected on the realignment parameters (Fair et al., 2019).

**Table 2.** Nuisance processes examined.

|  | SLFOs | Cardiac pulsatility | Breathing motion | Head motion |
| --- | --- | --- | --- | --- |
| **Recordings** | Pulse oximeter, breathing belt, fMRI global signal | Pulse oximeter | Breathing belt | fMRI |
| **Signals employed** | Heart rate, respiratory flow, global signal | Cardiac cycle | Breathing cycle | Realignment parameters and derivatives |
| **Model** | Kassinopoulos & Mitsis (2019a), *NeuroImage* | 3<sup>rd</sup> order RETROICOR<br>Glover et al. (2000),<br>*NeuroImage* | 3<sup>rd</sup> order RETROICOR<br>Glover et al. (2000),<br>*NeuroImage* | - |
| **Number of regressors** | 1 | 6 | 6 | 12 |

All nuisance regressors were high-pass filtered at 0.01 Hz to ensure similar spectral content with the fMRI data and thus avoid reintroduction of nuisance-related variation (Bright et al., 2017; Hallquist et al., 2013). The regressors were then normalized to zero mean and unit variance. **Supp. Fig. 9** demonstrates the spectral content of each nuisance process, as well as the effect of regressing out breathing motion from the realignment parameters.

### 5.4 Isolation of nuisance fluctuations from fMRI data

We propose a framework to isolate nuisance fluctuations for each of the aforementioned processes, which reflects the physiologically-driven fluctuations and head motion artifacts observed in the fMRI data (**Figure 7**). A similar methodology was used in (Bright and Murphy, 2015) to investigate whether preprocessing strategies remove variance associated to resting-state networks.

**Figure 7:**
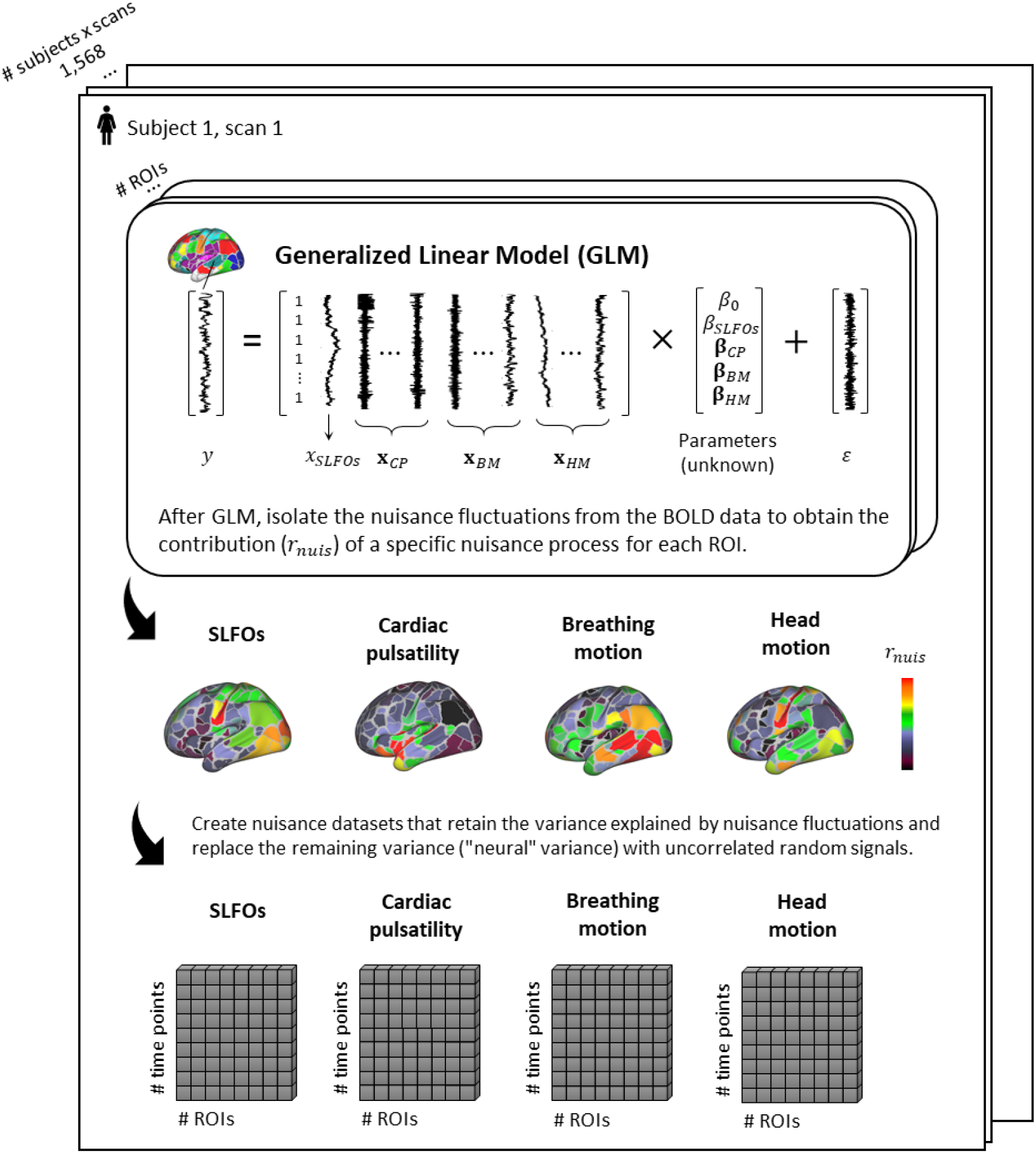
Graphical summary of the proposed framework for isolating the fluctuations for each physiological process. For each scan, the ROI time-series are modeled using the regressors related to systemic low frequency oscillations (SLFOs), cardiac pulsatility (CP), breathing motion (BM), and head motion (HM). Subsequently, the fraction of BOLD variance explained by each nuisance process is isolated and employed to generate synthetic datasets that only contain nuisance fluctuations.

Initially, the contribution of each nuisance process on the ROI time-series was quantified using a generalised linear model, formulated as:

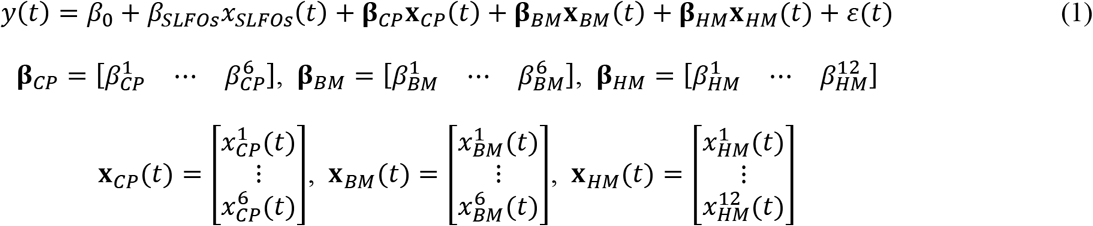

where *y* are ROI time-series from the minimally preprocessed data, *x_SLFOs_* is the physiological regressor modeling SLFOs, **x**_*CP*_(*t*) are the 6 physiological regressors modeling cardiac pulsatility, **x**_*BM*_(*t*) are the 6 physiological regressors modeling breathing motion, **x**_*HM*_(*t*) are the 12 regressors modeling head motion, {*β*_0_, *β_SLFOs_*, ***β**_CP_, **β**_BM_, **β**_HM_*} denote the parameters to be estimated, and *ε* is the error (or residual). As can be seen, all four nuisance processes (25 regressors in total) were included simultaneously in the regression to model the BOLD signal fluctuations in a specific ROI.

For each nuisance process, the estimated values 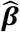 were multiplied by their corresponding regressors and added together to obtain the fluctuations of the nuisance process of interest (*ŷ_NPI_*(*t*)) within a specific ROI, and a “clean” time-series was calculated via removal of all other nuisance processes (*ŷ_NPI+Neur_*(*t*)), as follows:

SLFOs:

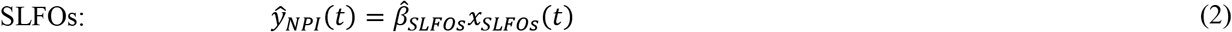

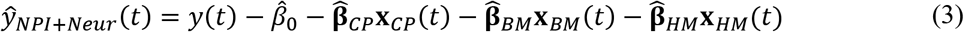

Cardiac pulsatility:

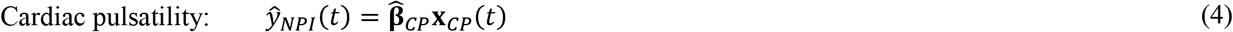

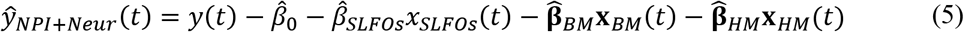

Breathing motion:

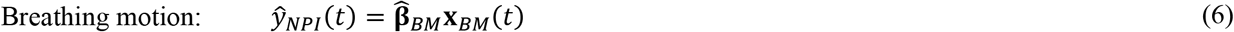

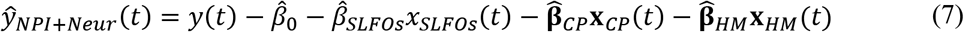

Head motion:

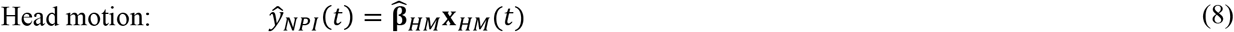

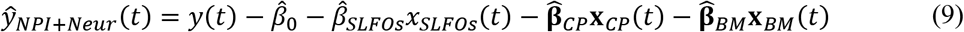

In this manner, we generated “cleaned” ROI time-series (*ŷ_NPI+Neur_*(*t*)) in which all considered noisy fluctuations were removed except the ones corresponding to the specific nuisance process being evaluated. The next step was to quantify the contribution of the latter to the remaining fluctuations within each ROI. To achieve this, the estimated nuisance signal was correlated to the “clean” ROI time-series:

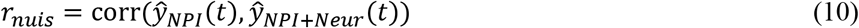

Subsequently, the estimated nuisance signal was subtracted from the “clean” ROI time-series to obtain what is typically considered the “neural” time-series. These time-series were correlated to the “clean” time-series to quantify the contribution of the “neural” variations to the total ROI signal fluctuations:

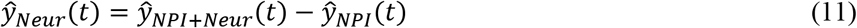

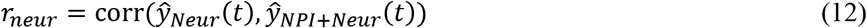

Afterwards, nuisance datasets for each process were created by scaling the estimated nuisance signal within each ROI with its corresponding correlation coefficient *r_nuis_* and adding Gaussian random signals (*ξ*(*t*)) scaled with *r_neur_*. This is expressed as:

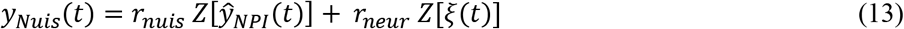

where *Z*[·] denotes normalization to zero mean and unit variance.

Thus, this framework generated four synthetic nuisance datasets that contained the isolated fluctuations from each of the nuisance processes evaluated. In a sense, the ROI time-series in each nuisance dataset are equivalent to the term *ŷ_NPI+Neur_*(*t*), with the “neural” fluctuations replaced by random signals. These time-series were used to characterize the connectome profile of the nuisance processes without the presence of neurally-related signals, while maintaining the noise-to-signal ratio between physiological/motion-related noise and “neural” signal intact. Note that if the nuisance datasets consisted solely of artifactual fluctuations without the random signals added, this would result in an overestimation of the correlation fraction attributed to the nuisance processes that was present in the experimental fMRI data. This can be easily understood in the case of two ROI time-series that are weakly driven by SLFOs. As the contribution of this nuisance process to the aggregate ROI time-series would be small, the parameter 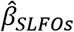 in Eq. 2 for both ROIs would be relatively small as well. However, without the addition of random signals, the correlation of the time-series *ŷ_NPI_* associated with these two ROIs would be 1.00 (or −1.00 depending on the signs of the corresponding beta parameters) as Pearson’s correlation is a metric that is blind to the variance of the signals, thereby overestimating the contribution of SLFOs to the FC between those ROIs.

### 5.5 Estimation of static and time-varying functional connectivity (FC)

Using the pre-processed fMRI data from each of the 12 pipelines (see section 5.2.2), henceforth called “neural” datasets, and the 4 nuisance datasets, we performed Pearson correlation analyses between brain regions over the whole scan (static FC) and within sliding windows (time-varying FC). To quantify time-varying FC, the entire scan was split up into 62 sliding windows of 43.2 sec (60 samples) duration, with 70% overlap in time. Subsequently, for each scan, we computed the functional connectivity dynamics (FCD) matrix (Hansen et al., 2015), which is a symmetric matrix in which the entries (*i,j*) correspond to the Pearson correlations between the upper triangular elements of the FC matrices in windows *i* and *j*. The size of the FCD matrix was *W*×*W*, where *W* is the number of windows (62). Thus, while the static FC matrix characterizes the spatial structure of resting activity, the FCD matrix captures the temporal evolution of connectome correlations.

The analyses resulted in 32 matrices for each subject (**Figure 8**): 4 static FC and 4 FCD nuisance matrices (one for each physiological process considered), as well as 12 static FC and 12 FCD “neural” matrices (one for each preprocessing strategy evaluated). To quantify the influence of the nuisance processes on static FC and FCD for each preprocessing strategy, similarities between pairs of nuisance and “neural” matrices were evaluated by correlating their upper triangular values. Note that for the FCD matrices, upper triangular elements corresponding to the correlation between overlapping windows were disregarded because of high correlation by design (see block diagonal in **Figure 8**). All correlation values were Fisher z transformed.

**Figure 8:**
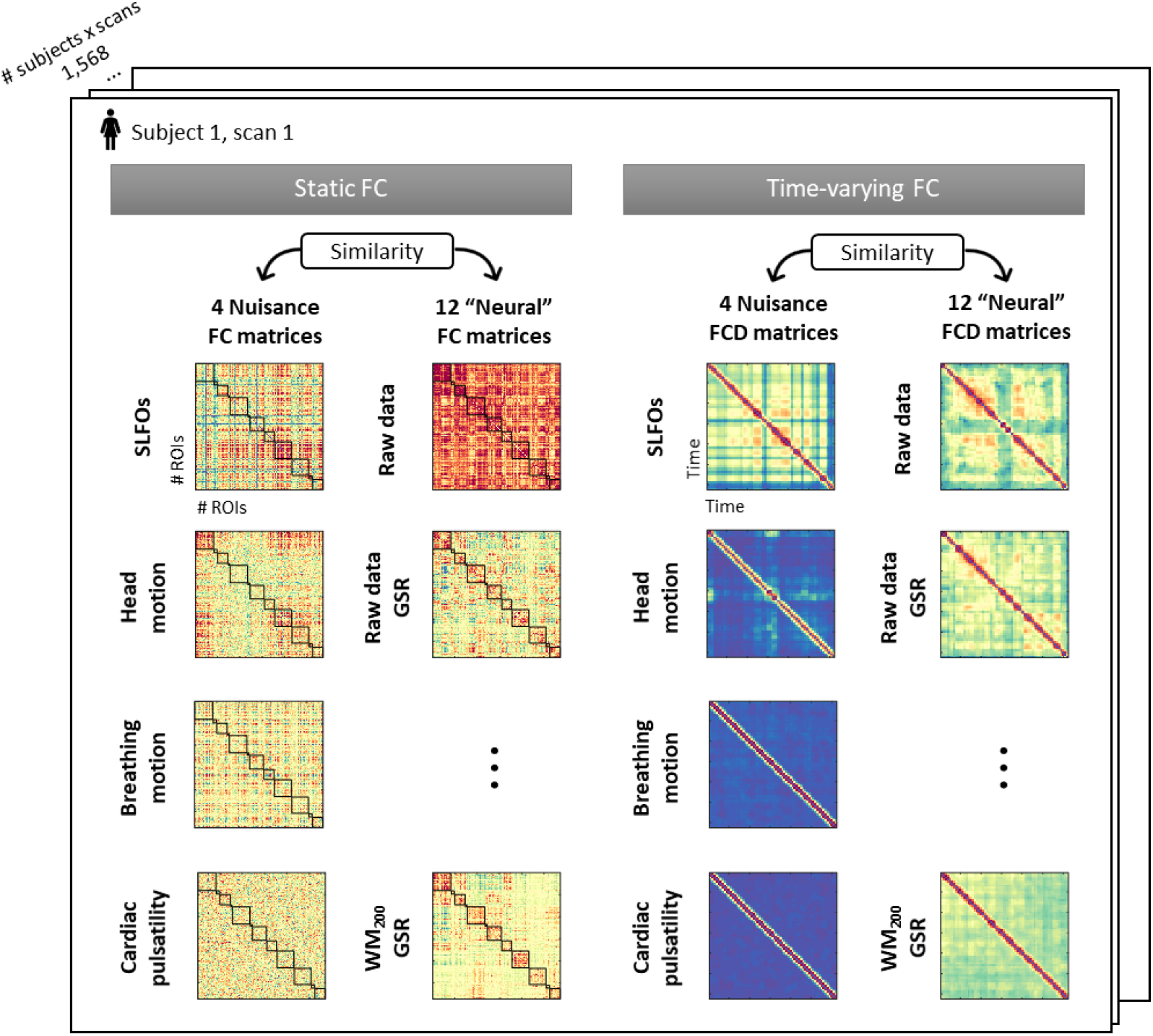
Illustration of the 32 connectivity matrices computed per scan. For both static and time-varying FC analyses, 4 nuisance connectivity matrices were computed using the generated nuisance datasets, as well as 12 “neural” connectivity matrices corresponding to the 12 pre-processing strategies evaluated (note that each of the 6 strategies listed in Table 1 was assessed with and without GSR). Static FC matrices were computed as the correlation across brain regions using the whole scan. Time-varying FC analysis constructed time-resolved connectivity matrices as the correlation between static FC matrices within sliding windows, known as functional connectivity dynamics (FCD) matrices (Hansen et al. 2015).

### 5.6 Connectome-based identification of individuals

We implemented a connectome-based identification of individual subjects using the static FC matrices to investigate the potential effect of “nuisance fingerprints” on the degree of subject specificity in individual FC metrics. The identification procedure, known as connectome fingerprinting, has been described in detail previously (Finn et al., 2015). Briefly, a database was first created consisting of all subjects’ FC matrices from a particular resting-state scan (**Supp. Fig. 10B**). An FC matrix from a specific subject and different resting-state scan was then selected and denoted as the target (*Subj_x_*). Pearson correlation coefficients (*r*_1_,…, *r_N_*) were computed between the upper triangular values of the target FC matrix and all the FC matrices in the database. If the highest correlation coefficient corresponded to a pair of FC matrices from the same subject, a successful identification was indicated (*ID_Subj_x__* = 1); otherwise, it was marked as an incorrect identification (*ID_Subj_x__* = 0). The identification test was repeated such that each subject serves as the target subject once, and then the ID values were averaged across subjects to obtain the identification accuracy of the database-target pair. This process was repeated until tests between all scanning sessions were performed. In total, 12 database-target combinations were computed (**Supp. Fig. 10A**). Identification was performed using the whole brain connectivity matrix, as well as based on edges from within and between networks. In the latter case, networks containing less than 10 ROIs were excluded from the analysis.

The connectome fingerprinting analysis was performed independently for each physiological process, as well as for each preprocessing strategy, using the generated FC matrices (**Figure 8**). This analysis was only performed on static FC matrices and not time-varying FC matrices because recurrent patterns of connectivity observed in the FCD matrices are not expected to occur at similar time instances between scans.

### 5.7 Statistics

To assess the significance of the results, surrogate nuisance datasets were generated via inter-subject surrogates (Lancaster *et al*., 2018), using fMRI data recorded from one subject’s scan and physiological signals recorded from a different subject’s scan (in the case of the head motion dataset, volume realignment parameters were employed). This procedure was performed for all 1,568 scans, creating a surrogate dataset for each of the four examined physiological processes with the same dimensions as the evaluated datasets. Note that when creating each surrogate dataset, only the nuisance process being examined is replaced by signals from a different subject, whereas all other nuisance regressors remain the same. The same analyses described before were repeated using these surrogate datasets, generating a chance distribution against which the results were compared. Thus, the significance of the contributions of each nuisance process to the BOLD signal fluctuations were tested against the contributions found using the surrogate data (two-sample t-test, *p* < 0.05, Bonferroni corrected), and the similarity between the nuisance and “neural” FC matrices was compared against the similarity obtained using surrogate nuisance FC matrices. Note that for visualization purposes, similarity values identified as outliers (> 3 SD) are not displayed in **Figure 2**.

To assess the significance of the fingerprinting analysis, we performed nonparametric permutation testing as in (Finn et al., 2015). Briefly, the described fingerprinting analysis was repeated for all scans and database-target combinations, but for the identification test, the subject identity in the database set was permuted. In this way, a “successful” identification was designated when the highest correlation coefficient was between the FC matrices of two different subjects. As expected, for all the nuisance and “neural” datasets, the identification accuracy estimated with nonparametric permutation testing was around 0.3%, which corresponds to the probability of selecting a specific subject from a group of 392 subjects when the subject is selected at random.

## Acknowledgments

This work was supported by the Natural Sciences and Engineering Research Council of Canada (Discovery Grant 34362 awarded to GDM), the Fonds de la Recherche du Quebec - Nature et Technologies (FRQNT; Team Grant PR191780-2016 awarded to GDM) and the Canada First Research Excellence Fund (awarded to McGill University for the Healthy Brains for Healthy Lives initiative). AXP and MK acknowledge funding from Québec Bio-imaging Network (QBIN). Data were provided by the Human Connectome Project, WU-Minn Consortium (Principal Investigators: David Van Essen and Kamil Ugurbil; 1U54MH091657) funded by the 16 NIH Institutes and Centers that support the NIH Blueprint for Neuroscience Research; and by the McDonnell Center for Systems Neuroscience at Washington University. We are extremely grateful to all Human Connectome Project study participants, who generously donated their time to make this resource possible.

## Author contributions

**Alba Xifra-Porxas**, Conceptualization, Methodology, Software, Formal analysis, Investigation, Data curation, Visualization, Writing–original draft, Writing–review & editing

**Michalis Kassinopoulos**, Conceptualization, Methodology, Investigation, Data curation, Writing–review & editing

**Georgios D. Mitsis**, Conceptualization, Supervision, Writing–review & editing

## Supplementary material

**Supp. Fig. 1.**
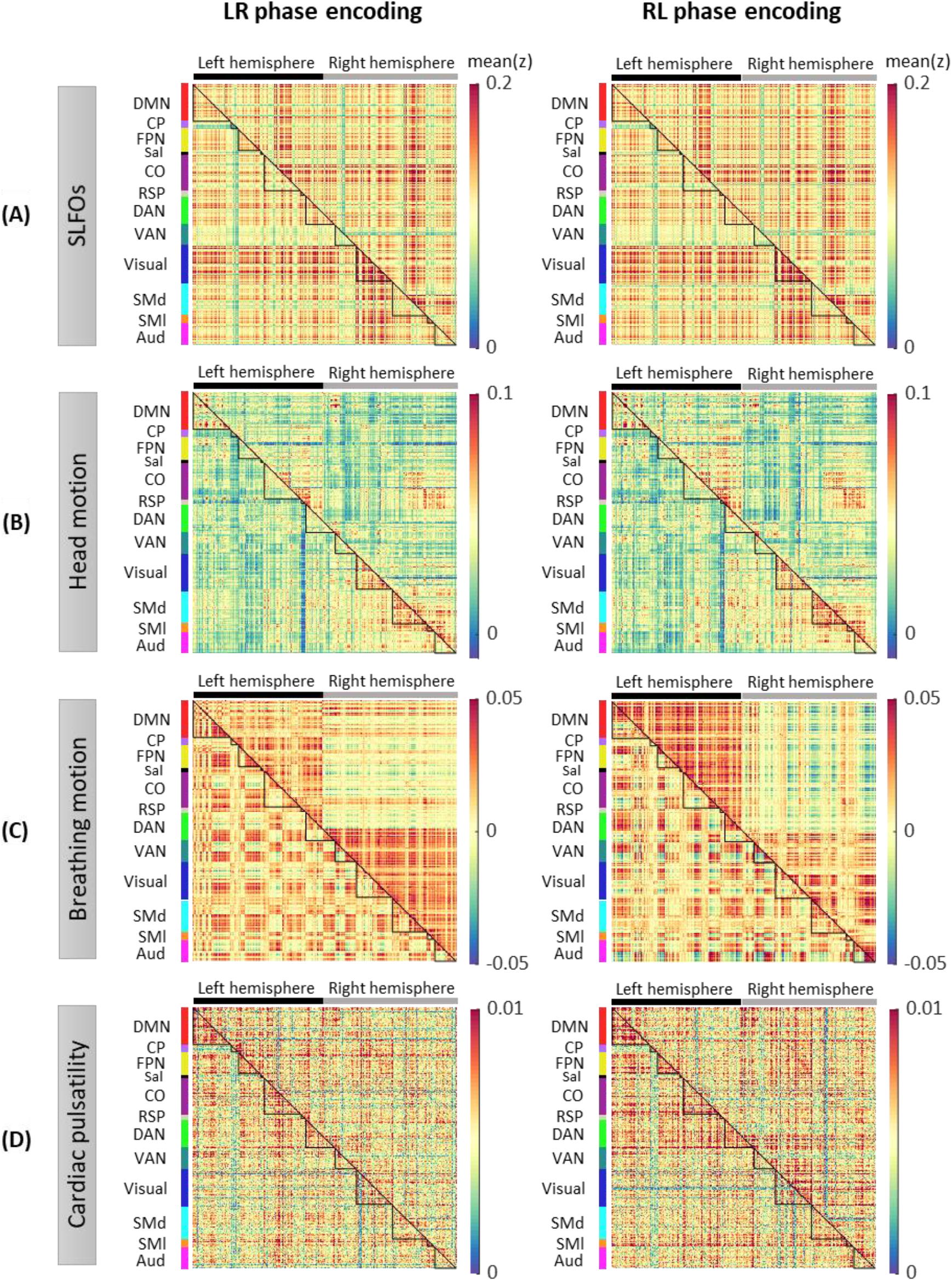
Whole-brain connectome patterns induced by nuisance processes, separately for scans with LR and RL phase encoding. Group averaged nuisance FC matrices across 784 scans based on their LR and RL phase encoding direction for (**A**) SLFOs, (**B**) head motion, (**C**) breathing motion, and (**D**) cardiac pulsatility.

**Supp. Fig. 2.**
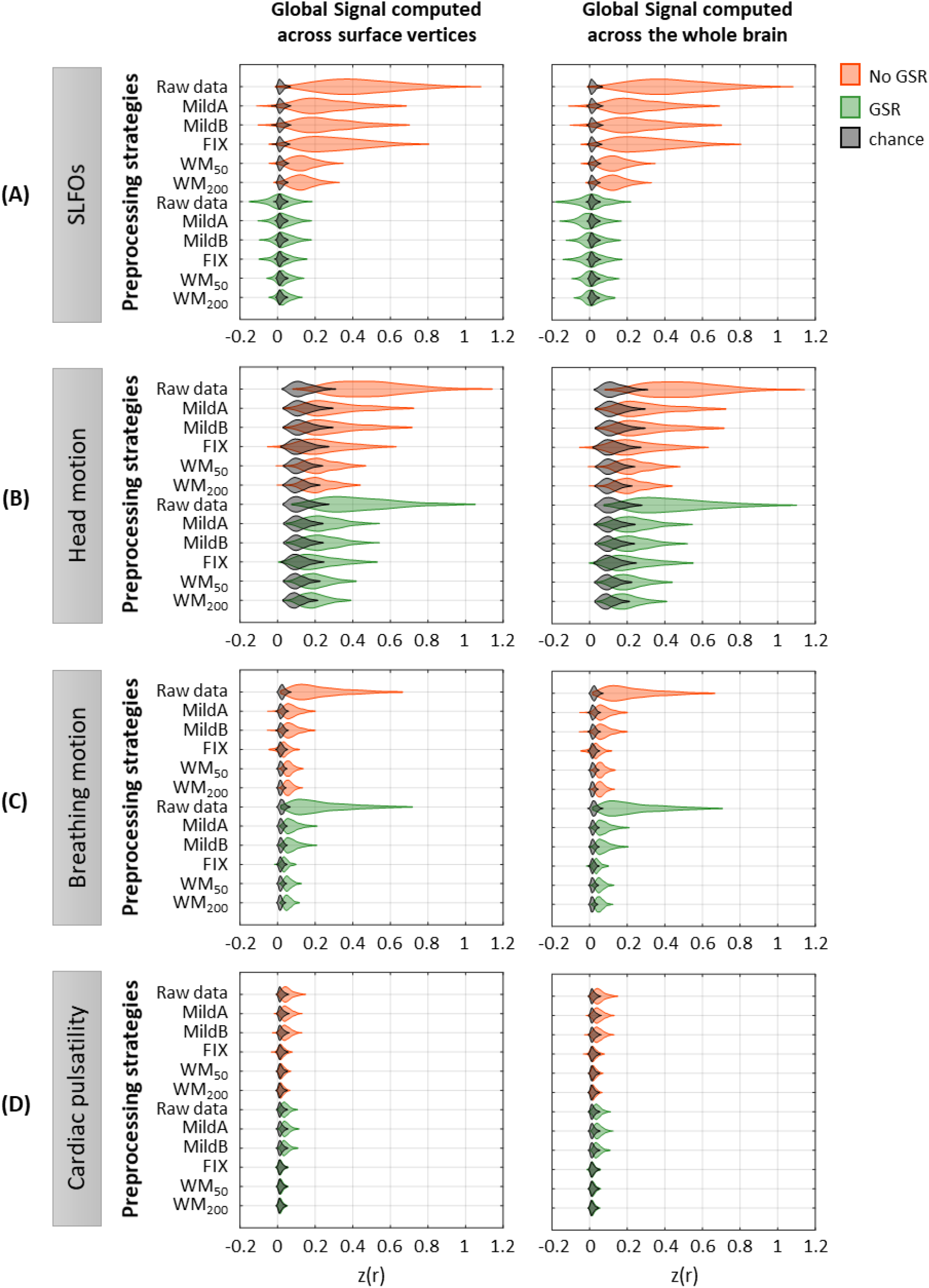
Effectiveness of preprocessing strategies in reducing the whole-brain connectivity profiles effects of each nuisance process for two different global signal calculation methods. Distribution of Pearson correlation coefficients across all 1,568 scans between the “neural” FC matrix after each preprocessing pipeline and nuisance FC matrices associated to (**A**) SLFOs, (**B**) head motion, (**C**) breathing motion, and (**D**) cardiac pulsatility, both when the global signal was computed across vertices in surface space (left column), and when the global signal was computed across the whole brain in volumetric space (right column). Correlation values were Fisher z transformed.

**Supp. Fig. 3.**
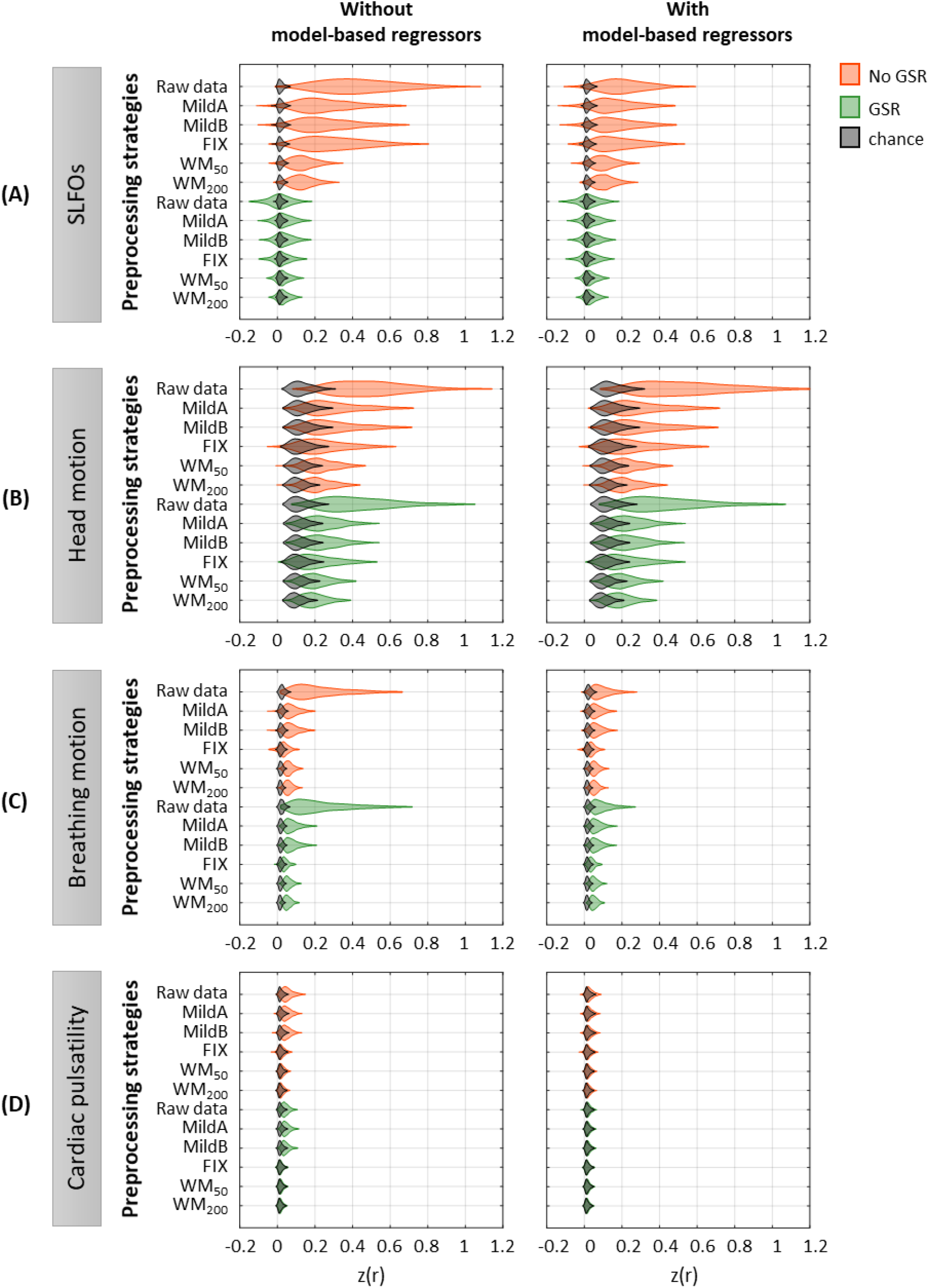
Effectiveness of preprocessing strategies in reducing the whole-brain connectivity effect of each nuisance process, with and without including model-based regressors. Distribution of Pearson correlation coefficients across all 1,568 runs between the “neural” FC matrix after each preprocessing pipeline and nuisance FC matrices associated to (**A**) SLFOs, (**B**) head motion, (**C**) breathing motion, and (**D**) cardiac pulsatility, when physiological regressors obtained from model-based techniques were included (right column) or not (left column) as nuisance regressors in the preprocessing strategies. Correlation values were Fisher z transformed.

**Supp. Fig. 4.**
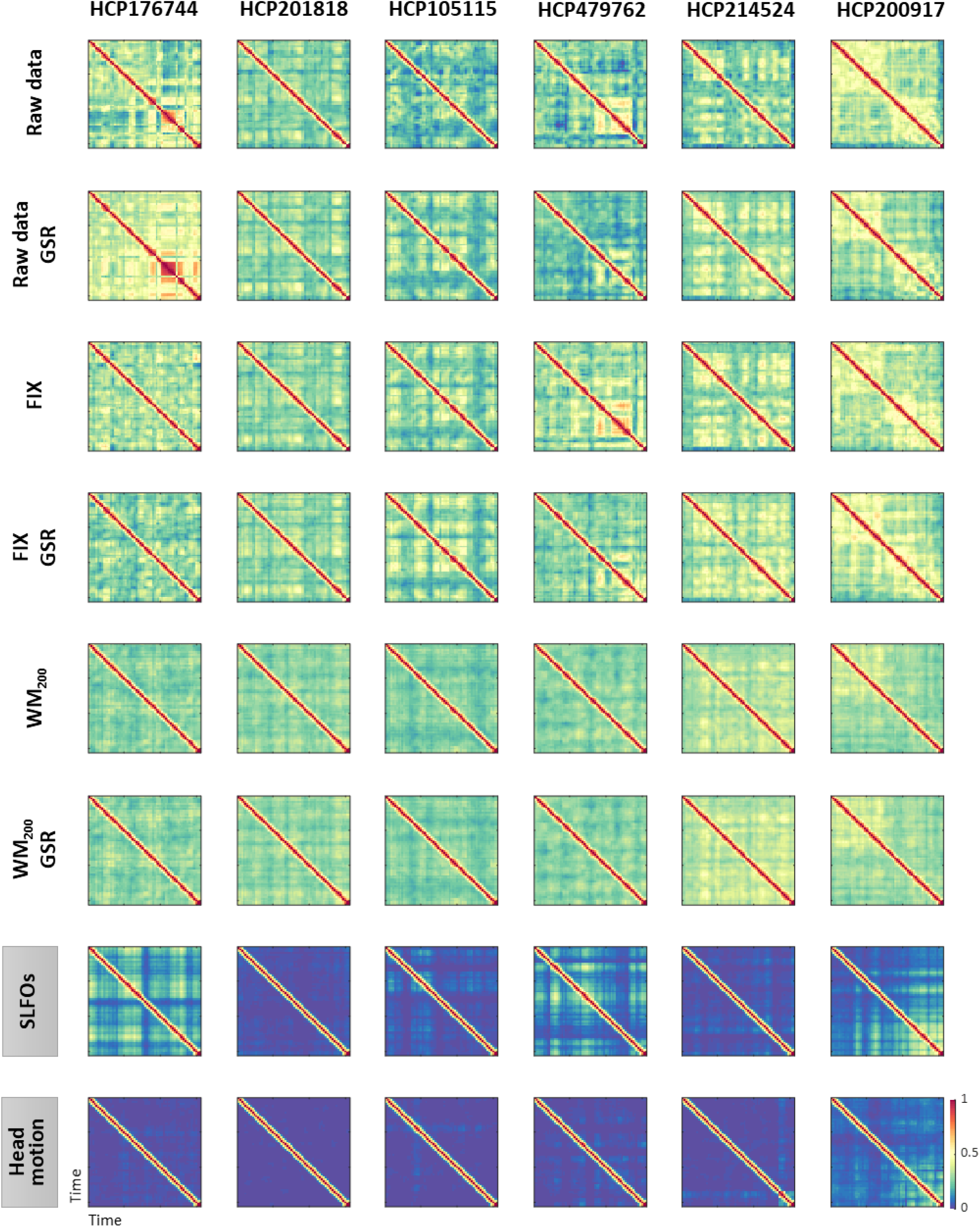
Examples of functional connectivity dynamics (FCD) profiles. Illustrative examples of FCD matrices from specific HCP subjects for several pre-processing pipelines (rows 1-6), SLFOs and head motion (rows 7 and 8, respectively). All the examples are from the HCP scan Rest1_LR. These subjects did not exhibit a large resemblance between FCD matrices computed from the “neural” datasets and FCD matrices computed from the nuisance datasets of SLFOs and head motion. Note that the WM_200_ preprocessing strategy substantially diminished the recurrent FC patterns.

**Supp. Fig. 5.**
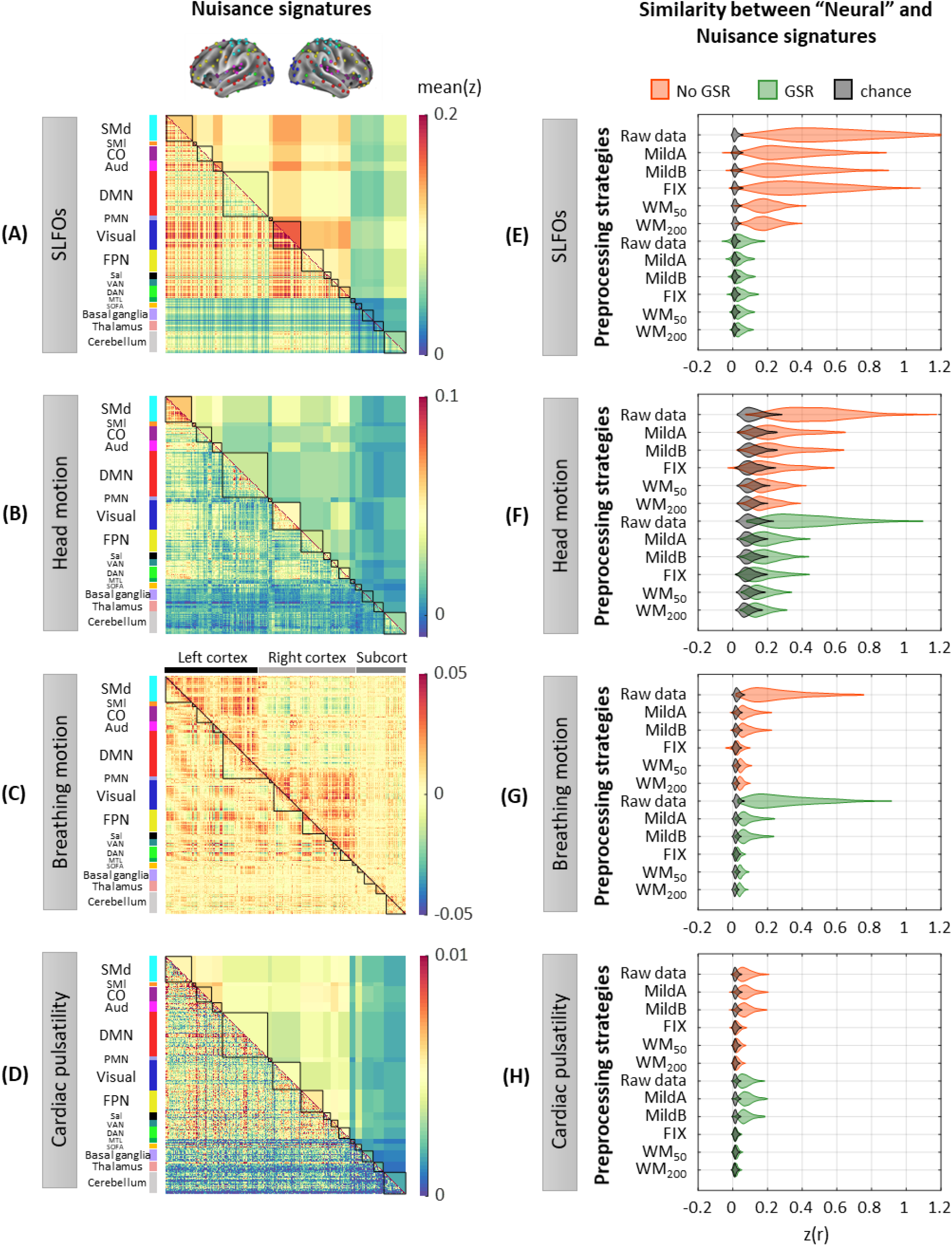
Whole-brain connectome patterns induced by nuisance processes and effect of preprocessing strategies, using the Seitzman atlas. (**A-D**) Group averaged “physiological” FC across all 1,568 scans for (**A**) SLFOs, (**B**) head motion, (**C**) breathing motion, and (**D**) cardiac pulsatility. (**E-H**) Distribution of Pearson correlation coefficients across all 1,568 scans between the “neural” FC matrix after each preprocessing pipeline and nuisance FC matrices associated to (**E**) SLFOs, (**F**) head motion, (**G**) breathing motion, and (**H**) cardiac pulsatility. Correlation values were Fisher z transformed.

**Supp. Fig. 6.**
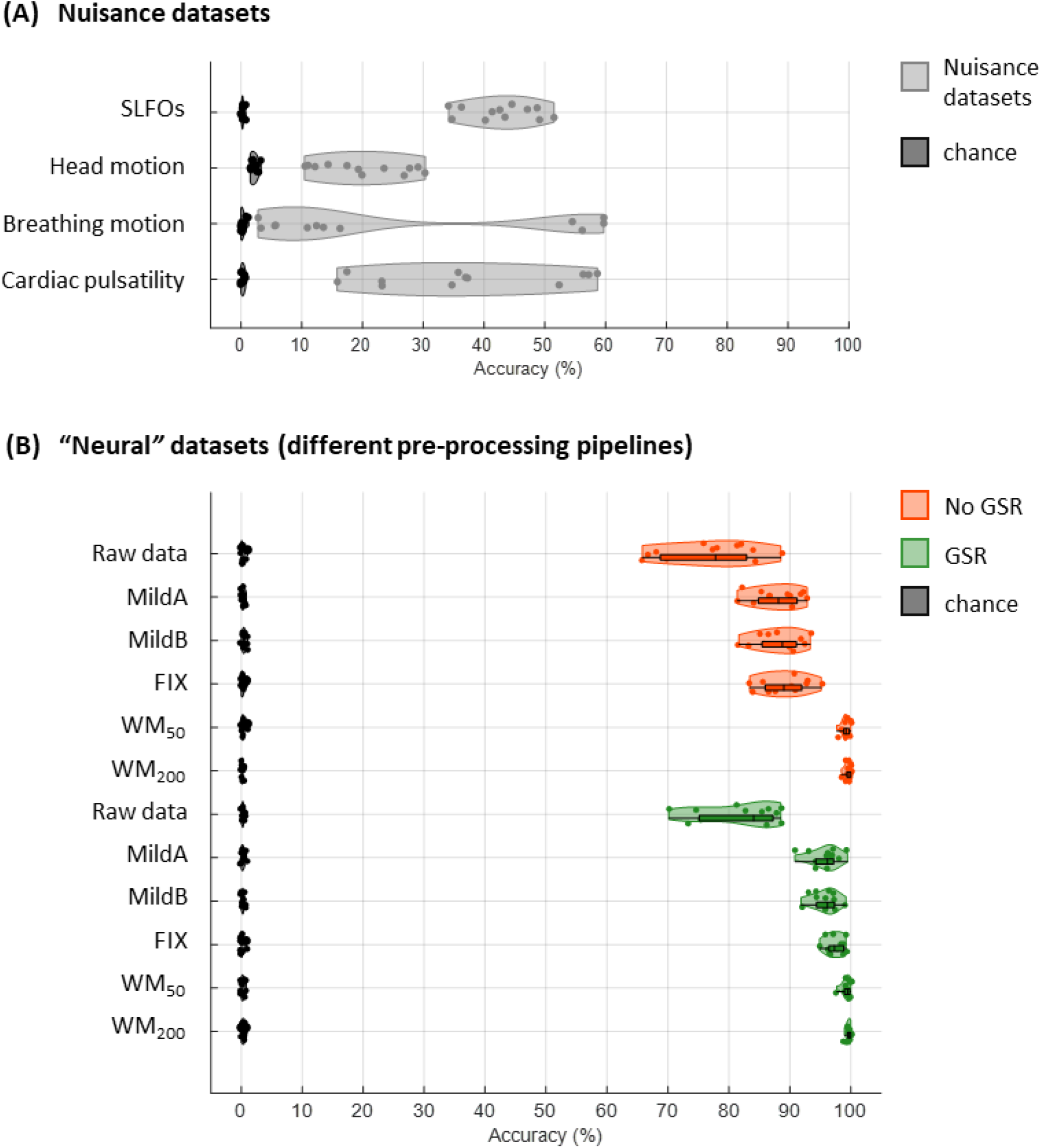
Connectome fingerprinting results for the Seitzman atlas. (**A**) Fingerprinting accuracy obtained using the static FC matrices from the generated nuisance datasets where non-neural fluctuations were isolated from the BOLD data. (**B**) Fingerprinting accuracy obtained using the static FC matrices generated from each of the preprocessing strategies evaluated.

**Supp. Fig. 7.**
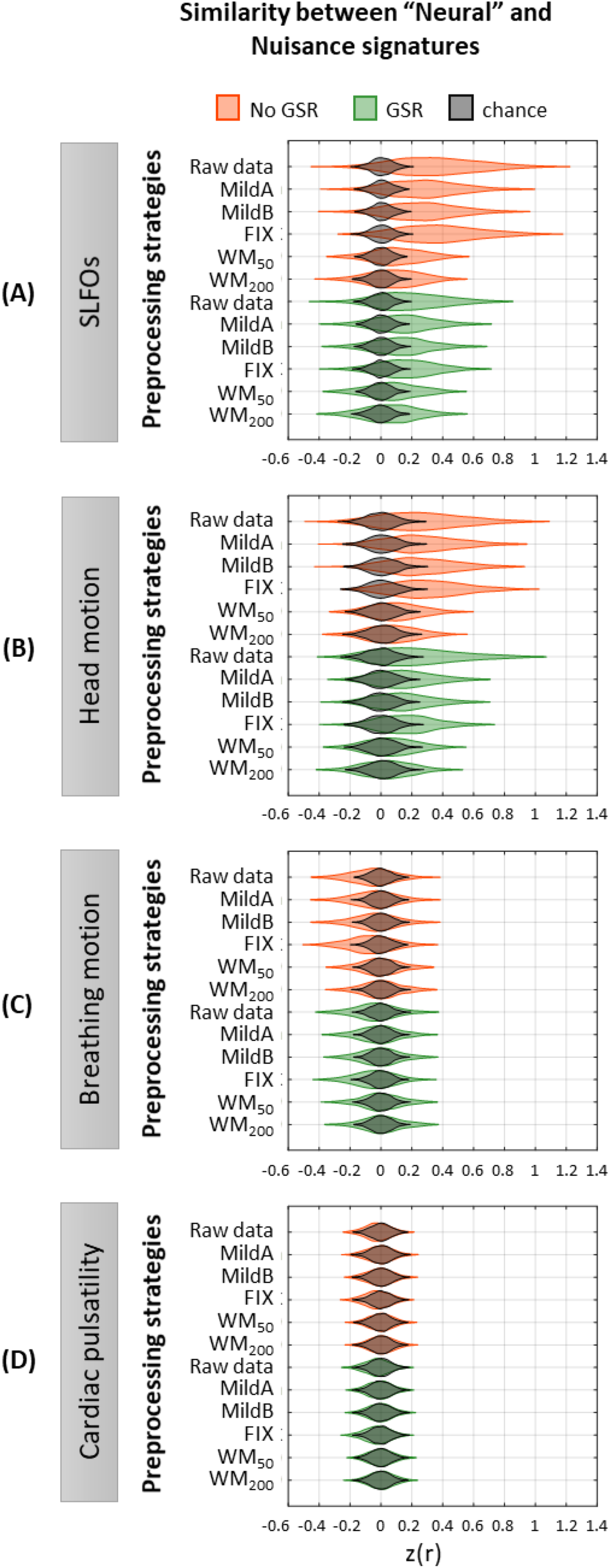
Effectiveness of preprocessing strategies in reducing functional connectivity dynamics (FCD) profiles induced by physiological and motion processes, using the Seitzman atlas. Distribution of Pearson correlation coefficients across all 1,568 scans between the “neural” FCD matrix after each preprocessing pipeline and nuisance FCD matrices associated to (**A**) SLFOs, (**B**) head motion, (**C**) breathing motion, and (**D**) cardiac pulsatility. Correlation values were Fisher z transformed.

**Supp. Fig. 8.**
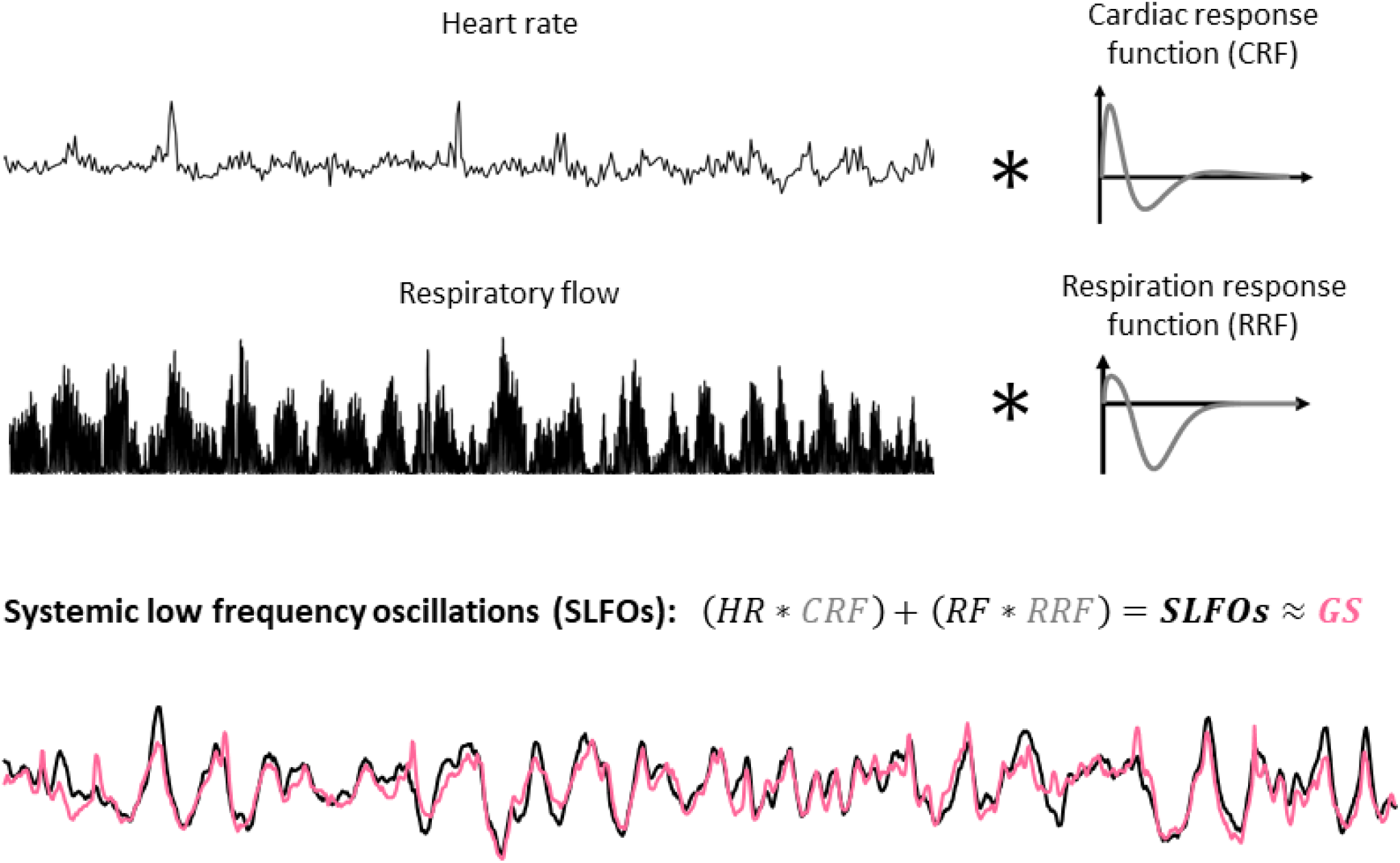
Illustration of methodology for modeling systemic low frequency oscillations (SLFOs) using the traces of heart rate and breathing activity. The respiratory flow was defined as the square of the derivative of the breathing signal. The cardiac and respiration response functions were estimated separately for each scan, using the methodology proposed in (Kassinopoulos and Mitsis, 2019a). Heart rate and respiratory flow were convolved with the cardiac and respiration response functions, respectively, in order to obtain the fluctuations of SLFOs. As can be seen in the example, SLFOs exhibit high correlation with the global signal (GS; the average correlation is 0.65 across all scans considered in this study).

**Supp. Fig. 9.**
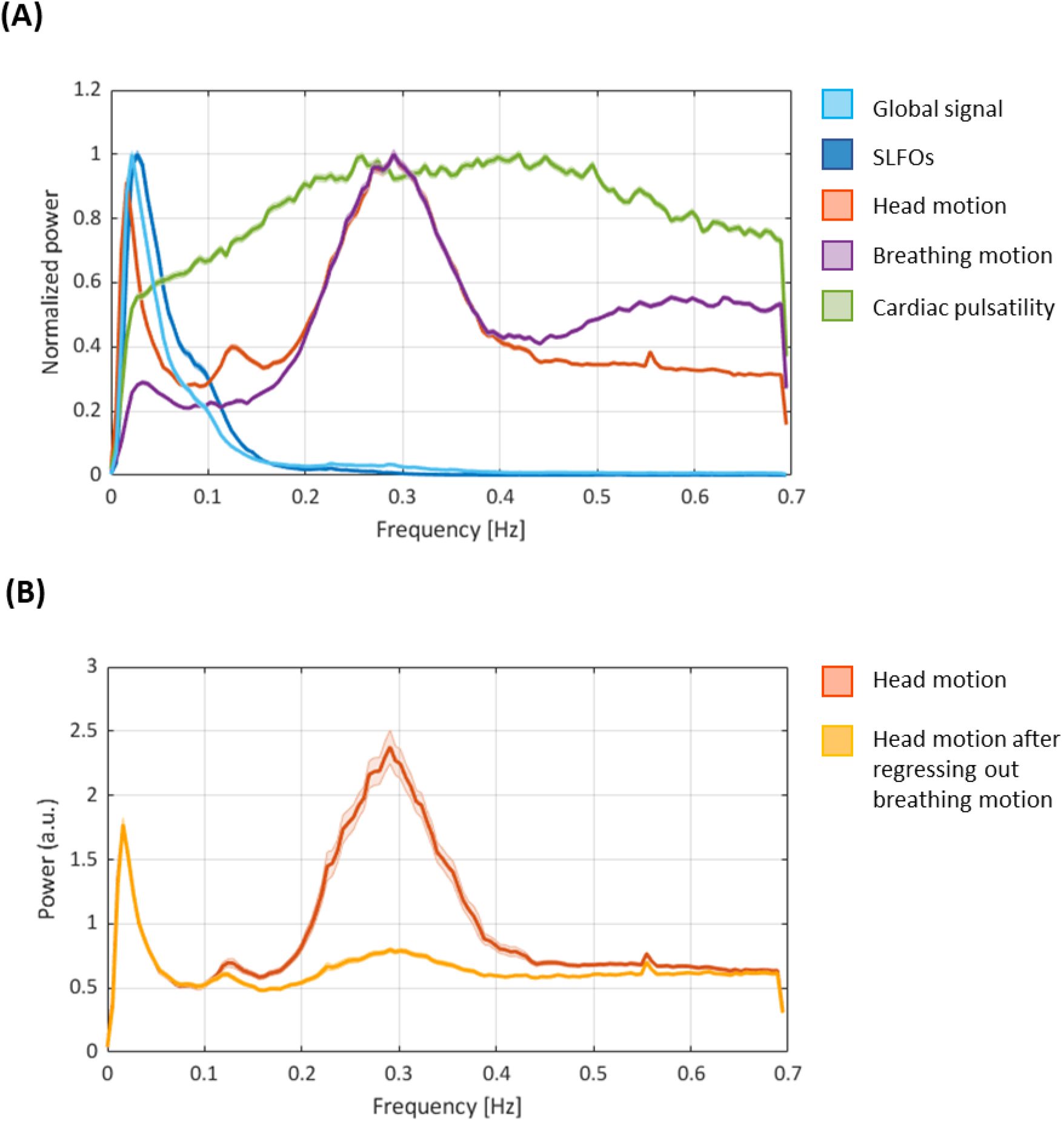
(A) Power spectral densities of the nuisance processes evaluated in the present study (as well as the global signal for reference). SLFOs mostly exhibit low-frequency fluctuations (<0.15 Hz). Breathing motion exhibits a peak at ~0.3 Hz, consistent with the average breathing rate across subjects. Head motion exhibits the same 0.3 Hz peak, underscoring the observation that realignment parameters are contaminated by breathing. We also observe a peak in head motion at ~0.12 Hz, recently attributed to deep breaths (Power et al. 2019), and a very narrow peak at 0.55 Hz that is likely due to scanner artifacts. The fMRI data is not sampled fast enough to capture cardiac pulsatility (~1 Hz), hence the effect of cardiac activity is aliased within a range of lower frequencies. Thus, any variation in heart rate during a scan as well as across subjects is likely to spread cardiac pulsatility artifacts across different frequencies, broadening the main spectral peak. (**B) Power spectral density of head motion before and after regressing out breathing motion.** Note the substantial decrease in power around 0.3 Hz after removing the effect of breathing motion. Both figures show the mean and standard error across subjects for scan Rest1_LR (similar results were observed for all other scans).

**Supp. Fig. 10.**
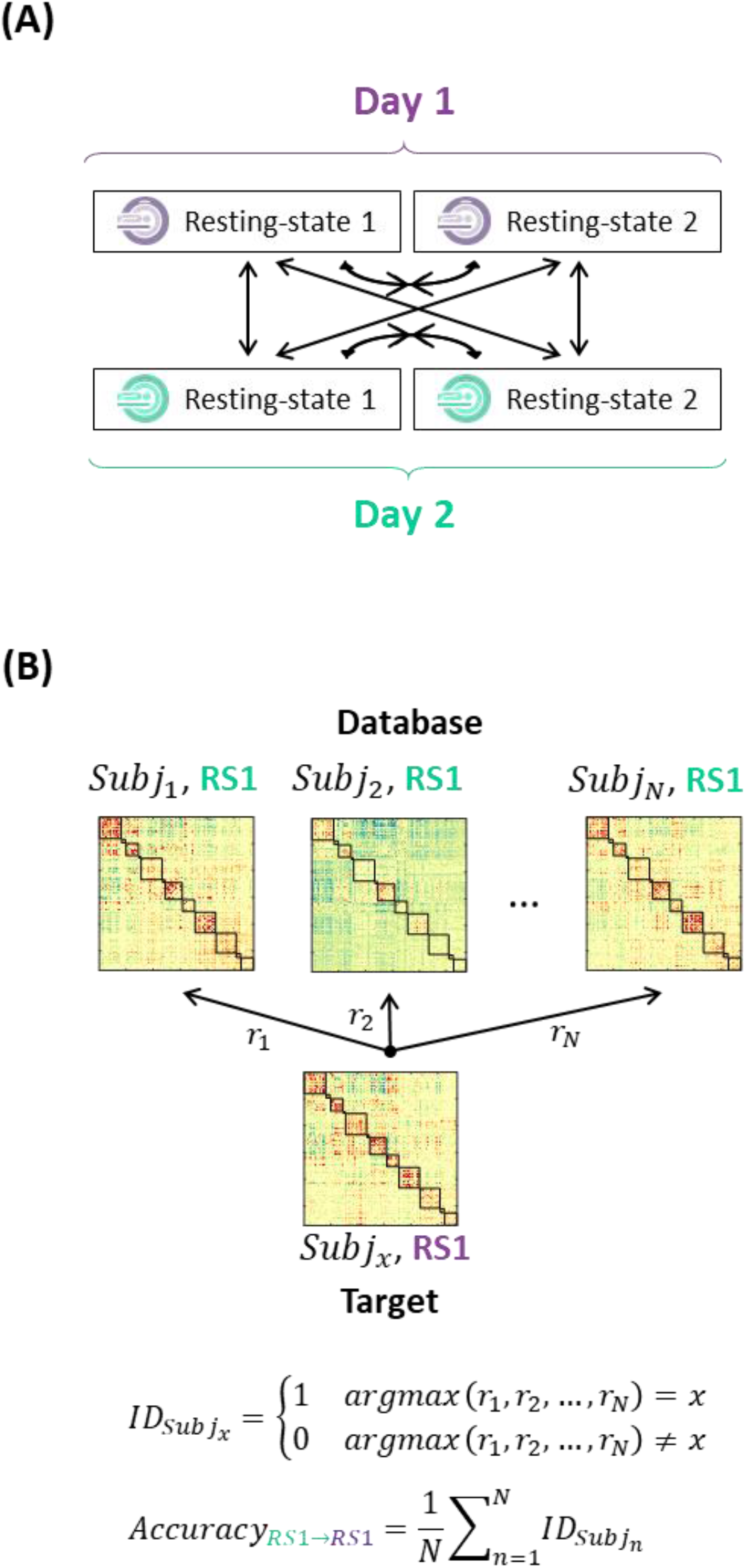
Connectome fingerprinting. (**A**) Diagram of the target-database combinations between resting-state scans. (**B**) Example on how to compute the identification accuracy for a target-database pair. Figure based on Finn et al. (2015).

## References

Alvares GA, Quintana DS, Hickie IB, Guastella AJ. 2016. Autonomic nervous system dysfunction in psychiatric disorders and the impact of psychotropic medications: A systematic review and meta-analysis. J Psychiatry Neurosci 41:89–104. doi:10.1503/jpn.140217

Aquino KM, Fulcher BD, Parkes L, Sabaroedin K, Fornito A. 2020. Identifying and removing widespread signal deflections from fMRI data: Rethinking the global signal regression problem. Neuroimage 212:116614. doi:10.1016/j.neuroimage.2020.116614

Baker AP, Brookes MJ, Rezek IA, Smith SM, Behrens T, Smith PJP, Woolrich M. 2014. Fast transient networks in spontaneous human brain activity. Elife 2014:1–18. doi:10.7554/eLife.01867

Baker JT, Holmes AJ, Masters GA, Yeo BTT, Krienen F, Buckner RL, Ongür D. 2014. Disruption of cortical association networks in schizophrenia and psychotic bipolar disorder. JAMA Psychiatry 71:109–118. doi:10.1001/jamapsychiatry.2013.3469

Batchvarov VN, Ghuran A, Smetana P, Hnatkova K, Harries M, Dilaveris P, Camm AJ, Malik M. 2002. QT-RR relationship in healthy subjects exhibits substantial intersubject variability and high intrasubject stability. Am J Physiol - Hear Circ Physiol 282:2356–2363. doi:10.1152/ajpheart.00860.2001

Battaglia D, Thomas B, Hansen EC, Chettouf S, Daffertshofer A, McIntosh AR, Zimmermann J, Ritter P, Jirsa V. 2017. Functional Connectivity Dynamics of the Resting State across the Human Adult Lifespan. bioRxiv 107243. doi:10.1101/107243

Benjamin BR, Valstad M, Elvsåshagen T, Jönsson EG, Moberget T, Winterton A, Haram M, Høegh MC, Lagerberg T V., Steen NE, Larsen L, Andreassen OA, Westlye LT, Quintana DS. 2020. Reduced heart rate variability is associated with disease severity in psychosis spectrum disorders. OSF Prepr. doi:10.31219/osf.io/upj3f

Bernier M, Cunnane SC, Whittingstall K. 2018. The morphology of the human cerebrovascular system. Hum Brain Mapp 1–14. doi:10.1002/hbm.24337

Bijsterbosch JD, Woolrich MW, Glasser MF, Robinson EC, Beckmann CF, Van Essen DC, Harrison SJ, Smith SM. 2018. The relationship between spatial configuration and functional connectivity of brain regions. Elife 7:1–27. doi:10.7554/eLife.32992

Billings JCW, Keilholz SD. 2018. The Not-So-Global BOLD Signal. Brain Connect 8:brain.2017.0517. doi:10.1089/brain.2017.0517

Birn RM, Cornejo MD aniel., Molloy EK, Patriat R, Meier TB, Kirk GR, Nair VA, Meyerand ME, Prabhakaran V. 2014. The influence of physiological noise correction on test-retest reliability of resting-state functional connectivity. Brain Connect 4:511–522. doi:10.1089/brain.2014.0284

Birn RM, Diamond JB, Smith MA, Bandettini PA. 2006. Separating respiratory-variation-related fluctuations from neuronalactivity-related fluctuations in fMRI. Neuroimage 31:1536–1548. doi:10.1016/j.neuroimage.2006.02.048

Birn RM, Murphy K, Bandettini PA. 2008a. The effect of respiration variations on independent component analysis results of resting state functional connectivity. Hum Brain Mapp 29:740–750. doi:10.1002/hbm.20577

Birn RM, Smith MA, Jones TB, Bandettini PA. 2008b. The respiration response function: The temporal dynamics of fMRI signal fluctuations related to changes in respiration. Neuroimage 40:644–654. doi:10.1016/j.neuroimage.2007.11.059

Biswal B, Yetkin FZ, Haughton VM, Hyde JS. 1995. Functional connectivity in the motor cortex of resting human brain using echo-planar MRI. Magn Reson Med. doi:10.1002/mrm.1910340409

Bright MG, Murphy K. 2015. Is fMRI “noise” really noise? Resting state nuisance regressors remove variance with network structure. Neuroimage 114:158–169. doi:10.1016/j.neuroimage.2015.03.070

Bright MG, Tench CR, Murphy K. 2017. Potential pitfalls when denoising resting state fMRI data using nuisance regression. Neuroimage 154:159–168. doi:10.1016/j.neuroimage.2016.12.027

Bright MG, Whittaker JR, Driver ID, Murphy K. 2018. Vascular physiology drives functional brain networks. bioRxiv 475491. doi:10.1101/475491

Brookes MJ, Hale JR, Zumer JM, Stevenson CM, Francis ST, Barnes GR, Owen JP, Morris PG, Nagarajan SS. 2011. Measuring functional connectivity using MEG: Methodology and comparison with fcMRI. Neuroimage 56:1082–1104. doi:10.1016/j.neuroimage.2011.02.054

Brosch JR, Talavage TM, Ulmer JL, Nyenhuis JA. 2002. Simulation of human respiration in fMRI with a mechanical model. IEEE Trans Biomed Eng 49:700–707. doi:10.1109/TBME.2002.1010854

Burgess GC, Kandala S, Nolan D, Laumann TO, Power JD, Adeyemo B, Harms MP, Petersen SE, Barch DM. 2016. Evaluation of Denoising Strategies to Address Motion-Correlated Artifacts in Resting-State Functional Magnetic Resonance Imaging Data from the Human Connectome Project. Brain Connect 6:669–680. doi:10.1089/brain.2016.0435

Byrge L, Kennedy DP. 2018. Identifying and characterizing systematic temporally-lagged BOLD artifacts. Neuroimage 171:376–392. doi:10.1016/j.neuroimage.2017.12.082

Caballero-Gaudes C, Reynolds RC. 2017. Methods for cleaning the BOLD fMRI signal. Neuroimage 154:128–149. doi:10.1016/j.neuroimage.2016.12.018

Carbonell F, Bellec P, Shmuel A. 2011. Global and System-Specific Resting-State fMRI Fluctuations Are Uncorrelated: Principal Component Analysis Reveals Anti-Correlated Networks. Brain Connect 1:496–510. doi:10.1089/brain.2011.0065

Chan MY, Park DC, Savalia NK, Petersen SE, Wig GS. 2014. Decreased segregation of brain systems across the healthy adult lifespan. Proc Natl Acad Sci 111:E4997–E5006. doi:10.1073/pnas.1415122111

Chang C, Cunningham JP, Glover GH. 2009. Influence of heart rate on the BOLD signal: The cardiac response function. Neuroimage 44:857–869. doi:10.1016/j.neuroimage.2008.09.029

Chang C, Glover GH. 2010. Time-frequency dynamics of resting-state brain connectivity measured with fMRI. Neuroimage 50:81–98. doi:10.1016/j.neuroimage.2009.12.011

Chang C, Glover GH. 2009a. Effects of model-based physiological noise correction on default mode network anti-correlations and correlations. Neuroimage 47:1448–1459. doi:10.1016/j.neuroimage.2009.05.012

Chang C, Glover GH. 2009b. Relationship between respiration, end-tidal CO2, and BOLD signals in resting-state fMRI. Neuroimage 47:1381–1393. doi:10.1016/j.neuroimage.2009.04.048

Chang C, Metzger CD, Glover GH, Duyn JH, Heinze HJ, Walter M. 2013. Association between heart rate variability and fluctuations in resting-state functional connectivity. Neuroimage 68:93–104. doi:10.1016/j.neuroimage.2012.11.038

Chen Heng, Nomi JS, Uddin LQ, Duan X, Chen Huafu. 2017. Intrinsic functional connectivity variance and state-specific under-connectivity in autism. Hum Brain Mapp 38:5740–5755. doi:10.1002/hbm.23764

Chen JE, Lewis LD, Chang C, Tian Q, Fultz NE, Ohringer NA, Rosen BR, Polimeni JR. 2020. Resting-state “physiological networks.” Neuroimage 213:116707. doi:10.1016/j.neuroimage.2020.116707

Chen JE, Polimeni JR, Bollmann S, Glover GH. 2019. On the analysis of rapidly sampled fMRI data. Neuroimage 188:807–820. doi:10.1016/j.neuroimage.2019.02.008

Ciric R, Wolf DH, Power JD, Roalf DR, Baum GL, Ruparel K, Shinohara RT, Elliott MA, Eickho SB, Davatzikos C, Gur RC, Gur RE, Bassett DS. 2017. Benchmarking of participant-level confound regression strategies for the control of motion artifact in studies of functional connectivity. Neuroimage 154:174–187. doi:10.1016/j.neuroimage.2017.03.020

Cole MW, Repovš G, Anticevic A. 2014. The frontoparietal control system: A central role in mental health. Neuroscientist 20:652–664. doi:10.1177/1073858414525995

Cole MW, Reynolds JR, Power JD, Repovs G, Anticevic A, Braver TS. 2013. Multi-task connectivity reveals flexible hubs for adaptive task control. Nat Neurosci 16:1348–1355. doi:10.1038/nn.3470

Collins CE, Airey DC, Young NA, Leitch DB, Kaas JH. 2010. Neuron densities vary across and within cortical areas in primates. Proc Natl Acad Sci U S A 107:15927–15932. doi:10.1073/pnas.1010356107

Dagli MS, Ingeholm JE, Haxby J V. 1999. Localization of cardiac-induced signal change in fMRI. Neuroimage 9:407–415. doi:10.1006/nimg.1998.0424

Damaraju E, Allen EA, Belger A, Ford JM, McEwen S, Mathalon DH, Mueller BA, Pearlson GD, Potkin SG, Preda A, Turner JA, Vaidya JG, Van Erp TG, Calhoun VD. 2014. Dynamic functional connectivity analysis reveals transient states of dysconnectivity in schizophrenia. NeuroImage Clin 5:298–308. doi:10.1016/j.nicl.2014.07.003

Demirtaş M, Tornador C, Falcón C, López-Solà M, Hernández-Ribas R, Pujol J, Menchón JM, Ritter P, Cardoner N, Soriano-Mas C, Deco G. 2016. Dynamic functional connectivity reveals altered variability in functional connectivity among patients with major depressive disorder. Hum Brain Mapp 37:2918–2930. doi:10.1002/hbm.23215

Dosenbach NUF, Fair DA, Miezin FM, Cohen AL, Wenger KK, Dosenbach RAT, Fox MD, Snyder AZ, Vincent JL, Raichle ME, Schlaggar BL, Petersen SE. 2007. Distinct brain networks for adaptive and stable task control in humans. Proc Natl Acad Sci 104:11073–11078. doi:10.1073/pnas.0704320104

Drysdale AT, Grosenick L, Downar J, Dunlop K, Mansouri F, Meng Y, Fetcho RN, Zebley B, Oathes DJ, Etkin A, Schatzberg AF, Sudheimer K, Keller J, Mayberg HS, Gunning FM, Alexopoulos GS, Fox MD, Pascual-Leone A, Voss HU, Casey BJ, Dubin MJ, Liston C. 2017. Resting-state connectivity biomarkers define neurophysiological subtypes of depression. Nat Med 23:28–38. doi:10.1038/nm.4246

Du Y, Pearlson GD, Yu Q, He H, Lin D, Sui J, Wu L, Calhoun VD. 2016. Interaction among subsystems within default mode network diminished in schizophrenia patients: A dynamic connectivity approach. Schizophr Res 170:55–65. doi:10.1016/j.schres.2015.11.021

Fair DA, Miranda-Dominguez O, Snyder AZ, Perrone A, Earl EA, Van AN, Koller JM, Feczko E, Tisdall MD, van der Kouwe A, Klein RL, Mirro AE, Hampton JM, Adeyemo B, Laumann TO, Gratton C, Greene DJ, Schlaggar BL, Hagler D, Watts R, Garavan H, Barch DM, Nigg JT, Petersen SE, Dale AM, Feldstein-Ewing SW, Nagel BJ, Dosenbach NUF. 2019. Correction of respiratory artifacts in MRI head motion estimates. Neuroimage 116400. doi:10.1016/j.neuroimage.2019.116400

Falahpour M, Refai H, Bodurka J. 2013. Subject specific BOLD fMRI respiratory and cardiac response functions obtained from global signal. Neuroimage 72:252–264. doi:10.1016/j.neuroimage.2013.01.050

Ferreira LK, Carolina A, Regina B, Kovacevic N, Morais G, Santos PP, Carneiro CDG, Kerr DS, Jr EA, Mcintosh AR, Busatto GF. 2016. Aging Effects on Whole-Brain Functional Connectivity in Adults Free of Cognitive and Psychiatric Disorders. Cereb Cortex 3851–3865. doi:10.1093/cercor/bhv190

Finn ES, Shen X, Scheinost D, Rosenberg MD, Huang J, Chun MM, Papademetris X, Constable RT. 2015. Functional connectome fingerprinting: Identifying individuals using patterns of brain connectivity. Nat Neurosci 18:1664–1671. doi:10.1038/nn.4135

Fox MD, Raichle ME. 2007. Spontaneous fluctuations in brain activity observed with functional magnetic resonance imaging. Nat Rev Neurosci 8:700–711. doi:10.1038/nrn2201

Friston KJ, Williams S, Howard R, Frackowiak RSJ, Turner R. 1996. Movement-related effects in fMRI time-series. Magn Reson Med 35:346–355. doi:10.1002/mrm.1910350312

Geerligs L, Renken RJ, Saliasi E, Maurits NM, Lorist MM. 2015. A Brain-Wide Study of Age-Related Changes in Functional Connectivity. Cereb Cortex 25:1987–1999. doi:10.1093/cercor/bhu012

Glasser MF, Coalson TS, Bijsterbosch JD, Harrison SJ, Harms MP, Anticevic A, Van Essen DC, Smith SM. 2018. Using temporal ICA to selectively remove global noise while preserving global signal in functional MRI data. Neuroimage 181:692–717. doi:10.1016/j.neuroimage.2018.04.076

Glasser MF, Smith SM, Marcus DS, Andersson JLR, Auerbach EJ, Behrens TEJ, Coalson TS, Harms MP, Jenkinson M, Moeller S, Robinson EC, Sotiropoulos SN, Xu J, Yacoub E, Ugurbil K, Van Essen DC. 2016. The Human Connectome Project’s neuroimaging approach. Nat Neurosci 19:1175–87. doi:10.1038/nn.4361

Glasser MF, Sotiropoulos SN, Wilson JA, Coalson TS, Fischl B, Andersson JL, Xu J, Jbabdi S, Webster M, Polimeni JR, Van Essen DC, Jenkinson M. 2013. The minimal preprocessing pipelines for the Human Connectome Project. Neuroimage 80:105–124. doi:10.1016/j.neuroimage.2013.04.127

Glover GH, Li T, Ress D. 2000. Image-Based Method for Retrospective Correction of Physiological Motion Effects in fMRI: RETROICOR. Magn Reson Med 44:162–167. doi:10.1002/1522-2594(200007)44:1<162::AID-MRM23>3.0.CO;2-E

Golestani AM, Chang C, Kwinta JB, Khatamian YB, Jean Chen J. 2015. Mapping the end-tidal CO2 response function in the resting-state BOLD fMRI signal: Spatial specificity, test-retest reliability and effect of fMRI sampling rate. Neuroimage 104:266–277. doi:10.1016/j.neuroimage.2014.10.031

Gonzalez-Castillo J, Caballero-Gaudes C, Topolski N, Handwerker DA, Pereira F, Bandettini PA. 2019. Imaging the spontaneous flow of thought: Distinct periods of cognition contribute to dynamic functional connectivity during rest. Neuroimage 202:116129. doi:10.1016/j.neuroimage.2019.116129

Gordon EM, Laumann TO, Adeyemo B, Huckins JF, Kelley WM, Petersen SE. 2016. Generation and Evaluation of a Cortical Area Parcellation from Resting-State Correlations. Cereb Cortex 26:288–303. doi:10.1093/cercor/bhu239

Gorgolewski KJ, Lurie D, Urchs S, Kipping JA, Craddock RC, Milham MP, Margulies DS, Smallwood J. 2014. A correspondence between individual differences in the brain’s intrinsic functional architecture and the content and form of self-generated thoughts. PLoS One 9. doi:10.1371/journal.pone.0097176

Gratton C, Dworetsky A, Coalson RS, Adeyemo B, Laumann TO, Wig GS, Kong TS, Gratton G, Fabiani M, Barch DM, Tranel D, Dominguez OM-, Fair DA, Dosenbach NUF, Snyder AZ, Perlmutter JS, Petersen SE, Campbell MC. 2020. Removal of high frequency contamination from motion estimates in single-band fMRI saves data without biasing functional connectivity. Neuroimage 116866. doi:10.1016/j.neuroimage.2020.116866

Gratton C, Koller JM, Shannon W, Greene DJ, Maiti B, Snyder AZ, Petersen SE, Perlmutter JS, Campbell MC. 2019. Emergent Functional Network Effects in Parkinson Disease. Cereb Cortex 29:2509–2523. doi:10.1093/cercor/bhy121

Gratton C, Laumann TO, Nielsen AN, Greene DJ, Gordon EM, Gilmore AW, Nelson SM, Coalson RS, Snyder AZ, Schlaggar BL, Dosenbach NUF, Petersen SE. 2018. Functional Brain Networks Are Dominated by Stable Group and Individual Factors, Not Cognitive or Daily Variation. Neuron 439–452. doi:10.1016/j.neuron.2018.03.035

Gratton C, Neta M, Sun H, Ploran EJ, Schlaggar BL, Wheeler ME, Petersen SE, Nelson SM. 2017. Distinct Stages of Moment-to-Moment Processing in the Cinguloopercular and Frontoparietal Networks. Cereb Cortex 27:2403–2417. doi:10.1093/cercor/bhw092

Griffanti L, Salimi-Khorshidi G, Beckmann CF, Auerbach EJ, Douaud G, Sexton CE, Zsoldos E, Ebmeier KP, Filippini N, Mackay CE, Moeller S, Xu J, Yacoub E, Baselli G, Ugurbil K, Miller KL, Smith SM. 2014. ICA-based artefact removal and accelerated fMRI acquisition for improved resting state network imaging. Neuroimage 95:232–247. doi:10.1016/j.neuroimage.2014.03.034

Hacker CD, Snyder AZ, Pahwa M, Corbetta M, Leuthardt EC. 2017. Frequency-specific electrophysiologic correlates of resting state fMRI networks. Neuroimage 149:446–457. doi:10.1016/j.neuroimage.2017.01.054

Hahamy A, Behrmann M, Malach R. 2015. The idiosyncratic brain: Distortion of spontaneous connectivity patterns in autism spectrum disorder. Nat Neurosci 18:302–309. doi:10.1038/nn.3919

Hallquist MN, Hwang K, Luna B. 2013. The nuisance of nuisance regression: Spectral misspecification in a common approach to resting-state fMRI preprocessing reintroduces noise and obscures functional connectivity. Neuroimage 82:208–225. doi:10.1016/j.neuroimage.2013.05.116

Hansen ECA, Battaglia D, Spiegler A, Deco G, Jirsa VK. 2015. Functional connectivity dynamics: Modeling the switching behavior of the resting state. Neuroimage 105:525–535. doi:10.1016/j.neuroimage.2014.11.001

He BJ, Snyder AZ, Zempel JM, Smyth MD, Raichle ME. 2008. Electrophysiological correlates of the brain’s intrinsic large-scale functional architecture. Proc Natl Acad Sci 105:16039–16044. doi:10.1073/pnas.0807010105

Hindriks R, Adhikari MH, Murayama Y, Ganzetti M, Mantini D, Logothetis NK, Deco G. 2016. Can sliding-window correlations reveal dynamic functional connectivity in resting-state fMRI? Neuroimage 127:242–256. doi:10.1016/j.neuroimage.2015.11.055

Hipp JF, Hawellek DJ, Corbetta M, Siegel M, Engel AK. 2012. Large-scale cortical correlation structure of spontaneous oscillatory activity. Nat Neurosci.

Hodkinson DJ, O’Daly O, Zunszain PA, Pariante CM, Lazurenko V, Zelaya FO, Howard MA, Williams SCR. 2014. Circadian and homeostatic modulation of functional connectivity and regional cerebral blood flow in humans under normal entrained conditions. J Cereb Blood Flow Metab 34:1493–1499. doi:10.1038/jcbfm.2014.109

Horien C, Shen X, Scheinost D, Constable RT. 2019. The individual functional connectome is unique and stable over months to years. Neuroimage 189:676–687. doi:10.1016/j.neuroimage.2019.02.002

Huck J, Wanner Y, Fan AP, Jäger AT, Grahl S, Schneider U, Villringer A, Steele CJ, Tardif CL, Bazin PL, Gauthier CJ. 2019. High resolution atlas of the venous brain vasculature from 7 T quantitative susceptibility maps. Brain Struct Funct 224:2467–2485. doi:10.1007/s00429-019-01919-4

Hunyadi B, Woolrich MW, Quinn AJ, Vidaurre D, De Vos M. 2019. A dynamic system of brain networks revealed by fast transient EEG fluctuations and their fMRI correlates. Neuroimage 185:72–82. doi:10.1016/j.neuroimage.2018.09.082

Hutchison RM, Womelsdorf T, Allen EA, Bandettini PA, Calhoun VD, Corbetta M, Della Penna S, Duyn JH, Glover GH, Gonzalez-Castillo J, Handwerker DA, Keilholz S, Kiviniemi V, Leopold DA, de Pasquale F, Sporns O, Walter M, Chang C. 2013. Dynamic functional connectivity: Promise, issues, and interpretations. Neuroimage 80:360–378. doi:10.1016/j.neuroimage.2013.05.079

Ji JL, Kulkarni K, Repovš G, Spronk M, Cole MW, Anticevic A. 2018. Mapping the human brain’s cortical-subcortical functional network organization. Neuroimage 185:35–57. doi:10.1016/j.neuroimage.2018.10.006

Jiang C, Yi L, Su S, Shi C, Long X, Xie G, Zhang L. 2016. Diurnal variations in neural activity of healthy human brain decoded with resting-state blood oxygen level dependent fMRI. Front Hum Neurosci 10:1–9. doi:10.3389/fnhum.2016.00634

Jo HJ, Saad ZS, Simmons WK, Milbury LA, Cox RW. 2010. Mapping sources of correlation in resting state FMRI, with artifact detection and removal. Neuroimage 52:571–582. doi:10.1016/j.neuroimage.2010.04.246

Kassinopoulos M, Mitsis GD. 2020. Physiological noise modeling in fMRI based on the pulsatile component of photoplethysmograph. bioRxiv. doi:10.1101/2020.06.01.128306

Kassinopoulos M, Mitsis GD. 2019a. White Matter Denoising Improves the Identifiability of Large-Scale Networks and Reduces the Effects of Motion in fMRI Functional Connectivity. bioRxiv.

Kassinopoulos M, Mitsis GD. 2019b. Identification of physiological response functions to correct for fluctuations in restingstate fMRI related to heart rate and respiration. Neuroimage 202. doi:10.1101/512855

Keller CJ, Bickel S, Honey CJ, Groppe DM, Entz L, Craddock RC, Lado FA, Kelly C, Milham M, Mehta AD. 2013. Neurophysiological investigation of spontaneous correlated and anticorrelated fluctuations of the BOLD signal. J Neurosci 33:6333–6342. doi:10.1523/JNEUROSCI.4837-12.2013

Kucyi A. 2018. Just a thought: How mind-wandering is represented in dynamic brain connectivity. Neuroimage 180:505–514. doi:10.1016/j.neuroimage.2017.07.001

Kucyi A, Davis KD. 2014. Dynamic functional connectivity of the default mode network tracks daydreaming. Neuroimage 100:471–480. doi:10.1016/j.neuroimage.2014.06.044

Kucyi A, Schrouff J, Bickel S, Foster BL, Shine JM, Parvizi J. 2018. Intracranial electrophysiology reveals reproducible intrinsic functional connectivity within human brain networks. J Neurosci 38:4230–4242. doi:10.1523/JNEUROSCI.0217-18.2018

Lancaster G, Iatsenko D, Pidde A, Ticcinelli V, Stefanovska A. 2018. Surrogate data for hypothesis testing of physical systems. Phys Rep 748:1–60. doi:10.1016/j.physrep.2018.06.001

Laumann TO, Snyder AZ, Mitra A, Gordon EM, Gratton C, Adeyemo B, Gilmore AW, Nelson SM, Berg JJ, Greene DJ, McCarthy JE, Tagliazucchi E, Laufs H, Schlaggar BL, Dosenbach NUF, Petersen SE. 2017. On the Stability of BOLD fMRI Correlations. Cereb Cortex 27:4719–4732. doi:10.1093/cercor/bhw265

Li J, Bolt T, Bzdok D, Nomi J, Yeo BT., Spreng RN, Uddin LQ. 2019a. Topography and behavioral relevance of the global signal in the human brain. Sci Rep 10:1–5. doi:10.1038/s41598-019-50750-8

Li J, Kong R, Liégeois R, Orban C, Tan Y, Sun N, Holmes AJ, Sabuncu MR, Ge T, Yeo BTT. 2019b. Global signal regression strengthens association between resting-state functional connectivity and behavior. Neuroimage 196:126–141. doi:10.1016/j.neuroimage.2019.04.016

Liu TT, Nalci A, Falahpour M. 2017. The global signal in fMRI: Nuisance or Information? Neuroimage 150:213–229. doi:10.1016/j.neuroimage.2017.02.036

Logothetis NK, Pauls J, Augath M, Trinath T, Oeltermann A. 2001. Neurophysiological investigation of the basis of the fMRI signal. Nature 412:150–157. doi:10.1038/35084005

Lurie DJ, Kessler D, Bassett DS, Betzel RF, Breakspear M, Keilholz S, Kucyi A, Liégeois R, Lindquist MA, McIntosh AR, Poldrack RA, Shine JM, Thompson WH, Bielczyk NZ, Douw20 L, Kraft D, Miller RL, Muthuraman M, Pasquini L, Razi A, Vidaurre D, Xie H, Calhoun VD. 2019. Questions and controversies in the study of time-varying functional connectivity in resting fMRI. Netw Neurosci. doi:https://doi.org/10.1162/netn_a_00116

Lydon-Staley DM, Ciric R, Satterthwaite TD, Basset DS. 2019. Evaluation of confound regression strategies for the mitigation of micromovement artifact in studies of dynamic resting-state functional connectivity and multilayer network modularity. Netw Neurosci 3:427–454. doi:10.1162/netn

Makowski C, Lepage M, Evans AC. 2019. Head motion: The dirty little secret of neuroimaging in psychiatry. J Psychiatry Neurosci 44:62–68. doi:10.1503/jpn.180022

Malik M, Hnatkova K, Sisakova M, Schmidt G. 2008. Subject-specific heart rate dependency of electrocardiographic QT, PQ, and QRS intervals. J Electrocardiol 41:491–497. doi:10.1016/j.jelectrocard.2008.06.022

Mash LE, Linke AC, Olson LA, Fishman I, Liu TT, Müller RA. 2019. Transient states of network connectivity are atypical in autism: A dynamic functional connectivity study. Hum Brain Mapp 40:2377–2389. doi:10.1002/hbm.24529

Matsui T, Murakami T, Ohki K. 2018. Neuronal Origin of the Temporal Dynamics of Spontaneous BOLD Activity Correlation. Cereb Cortex 1–13. doi:10.1093/cercor/bhy045

Mira-Dominguez O, Mills BD, Carpenter SD, Grant KA, Kroenke CD, Nigg JT, Fair DA. 2014. Connectotyping: Model based fingerprinting of the functional connectome. PLoS One 9. doi:10.1371/journal.pone.0111048

Morgan VL, Englot DJ, Rogers BP, Landman BA, Cakir A, Abou-Khalil BW, Anderson AW. 2017. Magnetic resonance imaging connectivity for the prediction of seizure outcome in temporal lobe epilepsy. Epilepsia 58:1251–1260. doi:10.1111/epi.13762

Mueller S, Wang D, Fox MD, Yeo BTT, Sepulcre J, Sabuncu MR, Shafee R, Lu J, Liu H. 2013. Individual Variability in Functional Connectivity Architecture of the Human Brain. Neuron 77:586–595. doi:10.1016/j.neuron.2012.12.028

Murphy K, Birn RM, Handwerker DA, Jones TB, Bandettini PA. 2009. The impact of global signal regression on resting state correlations: Are anti-correlated networks introduced? Neuroimage 44:893–905. doi:10.1016/j.neuroimage.2008.09.036

Murphy K, Fox MD. 2017. Towards a consensus regarding global signal regression for resting state functional connectivity MRI. Neuroimage 154:169–173. doi:10.1016/j.neuroimage.2016.11.052

Nalci A, Rao BD, Liu TT. 2019. Nuisance effects and the limitations of nuisance regression in dynamic functional connectivity fMRI. Neuroimage 184:1005–1031. doi:10.1016/j.neuroimage.2018.09.024

Nikolaou F, Orphanidou C, Papakyriakou P, Murphy K, Wise RG, Mitsis GD. 2016. Spontaneous physiological variability modulates dynamic functional connectivity in resting-state functional magnetic resonance imaging. Philos Trans R Soc A Math Phys Eng Sci 374:20150183. doi:10.1098/rsta.2015.0183

Noble S, Spann MN, Tokoglu F, Shen X, Constable RT, Scheinost D. 2017. Influences on the Test-Retest Reliability of Functional Connectivity MRI and its Relationship with Behavioral Utility. Cereb Cortex 27:5415–5429. doi:10.1093/cercor/bhx230

Noll DC, Schneider W. 1994. Theory, simulation, and compensation of physiological motion artifacts in functional MRI. Proc - Int Conf Image Process ICIP 3:40–44. doi:10.1109/ICIP.1994.413892

Ogawa S, Lee TM, Kay AR, Tank DW. 1990. Brain magnetic resonance imaging with contrast dependent on blood oxygenation. Proc Natl Acad Sci USA 87:9868–9872. doi:10.7566/JPSJ.84.064704

Orban C, Kong R, Li J, Chee MWL, Yeo BTT. 2020. Time of day is associated with paradoxical reductions in global signal fluctuation and functional connectivity. PLOS Biol 18:e3000602. doi:10.1371/journal.pbio.3000602

Özbay PS, Chang C, Picchioni D, Mandelkow H, Chappel-Farley MG, van Gelderen P, de Zwart JA, Duyn J. 2019. Sympathetic activity contributes to the fMRI signal. Commun Biol 2:1–9. doi:10.1038/s42003-019-0659-0

Parkes L, Fulcher B, Yücel M, Fornito A. 2018. An evaluation of the efficacy, reliability, and sensitivity of motion correction strategies for resting-state functional MRI. Neuroimage 171:415–436. doi:10.1016/j.neuroimage.2017.12.073

Patriat R, Molloy EK, Birn RM. 2015. Using Edge Voxel Information to Improve Motion Regression for rs-fMRI Connectivity Studies. Brain Connect 5:582–595. doi:10.1089/brain.2014.0321

Pinna GD, Maestri R, Torunski A, Danilowicz-Szymanowicz L, Szwoch M, La Rovere MT, Raczak G. 2007. Heart rate variability measures: A fresh look at reliability. Clin Sci 113:131–140. doi:10.1042/CS20070055

Pinto J, Nunes S, Bianciardi M, Dias A, Silveira LM, Wald LL, Figueiredo P. 2017. Improved 7 Tesla resting-state fMRI connectivity measurements by cluster-based modeling of respiratory volume and heart rate effects. Neuroimage 153:262–272. doi:10.1016/j.neuroimage.2017.04.009

Pitzalis MV, Mastropasqua F, Massari F, Forleo C, Di Maggio M, Passantino A, Colombo R, Di Biase M, Rizzon P. 1996. Short- and long-term reproducibility of time and frequency domain heart rate variability measurements in normal subjects. Cardiovasc Res 32:226–233. doi:10.1016/0008-6363(96)00086-7

Power JD. 2019. Temporal ICA has not properly separated global fMRI signals: A comment on Glasser et al. (2018). Neuroimage. doi:10.1016/j.neuroimage.2018.12.051

Power JD, Barnes KA, Snyder AZ, Schlaggar BL, Petersen SE. 2012. Spurious but systematic correlations in functional connectivity MRI networks arise from subject motion. Neuroimage 59:2142–2154. doi:10.1016/j.neuroimage.2011.10.018

Power JD, Lynch CJ, Dubin MJ, Silver BM, Martin A, Jones RM. 2020. Characteristics of respiratory measures in young adults scanned at rest, including systematic changes and “missed” deep breaths. Neuroimage 204:116234. doi:10.1016/j.neuroimage.2019.116234

Power JD, Mitra A, Laumann TO, Snyder AZ, Schlaggar BL, Petersen SE. 2014. Methods to detect, characterize, and remove motion artifact in resting state fMRI. Neuroimage 84:320–341. doi:10.1016/j.neuroimage.2013.08.048

Power JD, Plitt M, Gotts SJ, Kundu P, Voon V, Bandettini PA, Martin A. 2018. Ridding fMRI data of motion-related influences: Removal of signals with distinct spatial and physical bases in multiecho data. Proc Natl Acad Sci U S A 115:E2105–E2114. doi:10.1073/pnas.1720985115

Power JD, Plitt M, Laumann TO, Martin A. 2017. Sources and implications of whole-brain fMRI signals in humans. Neuroimage 146:609–625. doi:10.1016/j.neuroimage.2016.09.038

Power JD, Schlaggar BL, Petersen SE. 2015. Recent progress and outstanding issues in motion correction in resting state fMRI. Neuroimage 105:536–551. doi:10.1016/j.neuroimage.2014.10.044

Power JD, Silver BM, Dubin MJ, Martin A, Jones RM. 2019. Distinctions among real and apparent respiratory motions in human fMRI data. Neuroimage 201. doi:10.1101/601286

Prokopiou PC, Pattinson KTS, Wise RG, Mitsis GD. 2019. Modeling of dynamic cerebrovascular reactivity to spontaneous and externally induced CO2 fluctuations in the human brain using BOLD-fMRI. Neuroimage 186:533–548. doi:10.1016/j.neuroimage.2018.10.084

Pruim RHR, Mennes M, van Rooij D, Llera A, Buitelaar JK, Beckmann CF. 2015. ICA-AROMA: A robust ICA-based strategy for removing motion artifacts from fMRI data. Neuroimage 112:267–277. doi:10.1016/j.neuroimage.2015.02.064

Quaegebeur A, Lange C, Carmeliet P. 2011. The neurovascular link in health and disease: Molecular mechanisms and therapeutic implications. Neuron 71:406–424. doi:10.1016/j.neuron.2011.07.013

Raichle ME. 2015. The Brain’s Default Mode Network. Annu Rev Neurosci 38:433–447. doi:10.1146/annurev-neuro-071013-014030

Raj D, Anderson AW, Gore JC. 2001. Respiratory effects in human functional magnetic resonance imaging due to bulk susceptibility changes. Phys Med Biol 46:3331–3340. doi:10.1088/0031-9155/46/12/318

Raj D, Paey DP, Anderson AW, Kennan RP, Gore JC. 2000. A model for susceptibility artefacts from respiration in functional echo-planar magnetic resonance imaging. Phys Med Biol 45:3809–3820. doi:10.1088/0031-9155/45/12/321

Reland S, Ville NS, Wong S, Carrault G, Carré F. 2005. Reliability of heart rate variability in healthy older women at rest and during orthostatic testing. Aging Clin Exp Res 17:316–321. doi:10.1007/BF03324616

Robinson EC, Jbabdi S, Glasser MF, Andersson J, Burgess GC, Harms MP, Smith SM, Van Essen DC, Jenkinson M. 2014. MSM: A new flexible framework for multimodal surface matching. Neuroimage 100:414–426. doi:10.1016/j.neuroimage.2014.05.069

Sakoglu U, Calhoun VD, Pearlson GD, Kiehl KA, Wang YM, Michael AM. 2010. A method for evaluating dynamic functional network connectivity and task-modulation: Application to schizophrenia. Magn Reson Mater Physics, Biol Med. doi:10.1007/s10334-010-0197-8

Sala-Llonch R, Bartrés-Faz D, Junqué C. 2015. Reorganization of brain networks in aging: a review of functional connectivity studies. Front Psychol 6:663. doi:10.3389/fpsyg.2015.00663

Salimi-Khorshidi G, Douaud G, Beckmann CF, Glasser MF, Griffanti L, Smith SM. 2014. Automatic denoising of functional MRI data: Combining independent component analysis and hierarchical fusion of classifiers. Neuroimage 90:449–468. doi:10.1016/j.neuroimage.2013.11.046

Satterthwaite TD, Elliott MA, Gerraty RT, Ruparel K, Loughead J, Calkins ME, Eickhoff SB, Hakonarson H, Gur RC, Gur RE, Wolf DH. 2013. An improved framework for confound regression and filtering for control of motion artifact in the preprocessing of resting-state functional connectivity data. Neuroimage 64:240–256. doi:10.1016/j.neuroimage.2012.08.052

Satterthwaite TD, Wolf DH, Loughead J, Ruparel K, Elliott MA, Hakonarson H, Gur RC, Gur RE. 2012. Impact of in-scanner head motion on multiple measures of functional connectivity: Relevance for studies of neurodevelopment in youth. Neuroimage 60:623–632. doi:10.1016/j.neuroimage.2011.12.063

Savva AD, Kassinopoulos M, Smyrnis N, Matsopoulos GK, Mitsis GD. 2019. Effects of Motion Related Outliers in Dynamic Functional Connectivity Using the Sliding Window Method. J Neurosci Methods 108519. doi:10.1016/j.jneumeth.2019.108519

Schölvinck ML, Maier A, Ye FQ, Duyn JH, Leopold DA. 2010. Neural basis of global resting-state fMRI activity. Proc Natl Acad Sci USA 107:10238–43. doi:10.1073/pnas.0913110107

Seitzman BA, Gratton C, Laumann TO, Gordon EM, Adeyemo B, Dworetsky A, Kraus BT, Gilmore AW, Berg JJ, Ortega M, Nguyen A, Greene DJ, McDermott KB, Nelson SM, Lessov-Schlaggar CN, Schlaggar BL, Dosenbach NUF, Petersen SE. 2019a. Trait-like variants in human functional brain networks. Proc Natl Acad Sci. doi:10.1073/pnas.1902932116

Seitzman BA, Gratton C, Marek S, Raut R V., Dosenbach NUF, Schlaggar BL, Petersen SE, Greene DJ. 2019b. A set of functionally-defined brain regions with improved representation of the subcortex and cerebellum. Neuroimage 116290. doi:10.1016/j.neuroimage.2019.116290

Shannon BJ, Dosenbach RA, Su Y, Vlassenko AG, Larson-Prior LJ, Nolan TS, Snyder AZ, Raichle ME. 2013. Morning-evening variation in human brain metabolism and memory circuits. J Neurophysiol 109:1444–1456. doi:10.1152/jn.00651.2012

Shmuel A, Leopold DA. 2008. Neuronal correlates of spontaneous fluctuations in fMRI signals in monkey visual cortex: Implications for functional connectivity at rest. Hum Brain Mapp 29:751–761. doi:10.1002/hbm.20580

Shmueli K, van Gelderen P, de Zwart JA, Horovitz SG, Fukunaga M, Jansma JM, Duyn JH. 2007. Low-frequency fluctuations in the cardiac rate as a source of variance in the resting-state fMRI BOLD signal. Neuroimage 38:306–320. doi:10.1016/j.neuroimage.2007.07.037

Smith SM, Beckmann CF, Andersson J, Auerbach EJ, Bijsterbosch J, Douaud G, Duff E, Feinberg DA, Griffanti L, Harms MP, Kelly M, Laumann T, Miller KL, Moeller S, Petersen S, Power J, Salimi-Khorshidi G, Snyder AZ, Vu AT, Woolrich MW, Xu J, Yacoub E, Uğurbil K, Van Essen DC, Glasser MF. 2013. Resting-state fMRI in the Human Connectome Project. Neuroimage 80:144–168. doi:10.1016/j.neuroimage.2013.05.039

Smith SM, Fox PT, Miller KL, Glahn DC, Fox PM, Mackay CE, Filippini N, Watkins KE, Toro R, Laird AR, Beckmann CF. 2009. Correspondence of the brain’s functional architecture during activation and rest. Proc Natl Acad Sci USA 106:13040–5. doi:10.1073/pnas.0905267106

Tagliazucchi E, Laufs H. 2014. Decoding Wakefulness Levels from Typical fMRI Resting-State Data Reveals Reliable Drifts between Wakefulness and Sleep. Neuron 82:695–708. doi:10.1016/j.neuron.2014.03.020

Thompson GJ, Magnuson ME, Merritt MD, Schwarb H, Pan WJ, Mckinley A, Tripp LD, Schumacher EH, Keilholz SD. 2013a. Short-time windows of correlation between large-scale functional brain networks predict vigilance intraindividually and interindividually. Hum Brain Mapp 34:3280–3298. doi:10.1002/hbm.22140

Thompson GJ, Merritt MD, Pan WJ, Magnuson ME, Grooms JK, Jaeger D, Keilholz SD. 2013b. Neural correlates of time-varying functional connectivity in the rat. Neuroimage 83:826–836. doi:10.1016/j.neuroimage.2013.07.036

Tong Y, Frederick B de B. 2014. Studying the spatial distribution of physiological effects on BOLD signals using ultrafast fMRI. Front Hum Neurosci 8:1–8. doi:10.3389/fnhum.2014.00196

Tong Y, Hocke LM, Frederick BB. 2019. Low Frequency Systemic Hemodynamic “Noise” in Resting State BOLD fMRI: Characteristics, Causes, Implications, Mitigation Strategies, and Applications. Front Neurosci 13. doi:10.3389/fnins.2019.00787

Tong Y, Hocke LM, Nickerson LD, Licata SC, Lindsey KP, Frederick B de B. 2013. Evaluating the effects of systemic low frequency oscillations measured in the periphery on the independent component analysis results of resting state networks. Neuroimage 76:202–215. doi:10.1016/j.neuroimage.2013.03.019

Valenza G, Sclocco R, Duggento A, Passamonti L, Napadow V, Barbieri R, Toschi N. 2019. The central autonomic network at rest: Uncovering functional MRI correlates of time-varying autonomic outflow. Neuroimage. doi:10.1016/j.neuroimage.2019.04.075

Van de Moortele PF, Pfeuffer J, Glover GH, Ugurbil K, Hu X. 2002. Respiration-induced Bo fluctuations and their spatial distribution in the human brain at 7 Tesla. Magn Reson Med 47:888–895. doi:10.1002/mrm.10145

van Dijk KRA, Sabuncu MR, Buckner RL. 2012. The influence of head motion on intrinsic functional connectivity MRI. Neuroimage 59:431–438. doi:10.1016/j.neuroimage.2011.07.044

Van Essen DC, Smith SM, Barch DM, Behrens TEJ, Yacoub E, Ugurbil K. 2013. The WU-Minn Human Connectome Project: An overview. Neuroimage 80:62–79. doi:10.1016/j.neuroimage.2013.05.041

Van Essen DC, Ugurbil K, Auerbach E, Barch D, Behrens TEJ, Bucholz R, Chang A, Chen L, Corbetta M, Curtiss SW, Della Penna S, Feinberg D, Glasser MF, Harel N, Heath AC, Larson-Prior L, Marcus D, Michalareas G, Moeller S, Oostenveld R, Petersen SE, Prior F, Schlaggar BL, Smith SM, Snyder AZ, Xu J, Yacoub E. 2012. The Human Connectome Project: A data acquisition perspective. Neuroimage 62:2222–2231. doi:10.1016/j.neuroimage.2012.02.018

Vanderwal T, Eilbott J, Finn ES, Craddock RC, Turnbull A, Castellanos FX. 2017. Individual differences in functional connectivity during naturalistic viewing conditions. Neuroimage 157:521–530. doi:10.1016/j.neuroimage.2017.06.027

Vidaurre D, Hunt LT, Quinn AJ, Hunt BAE, Brookes MJ, Nobre AC, Woolrich MW. 2018. Spontaneous cortical activity transiently organises into frequency specific phase-coupling networks. Nat Commun 9. doi:10.1038/s41467-018-05316-z

Vigneau-Roy N, Bernier M, Descoteaux M, Whittingstall K. 2014. Regional variations in vascular density correlate with resting-state and task-evoked blood oxygen level-dependent signal amplitude. Hum Brain Mapp 35:1906–1920. doi:10.1002/hbm.22301

Wang C, Ong JL, Patanaik A, Zhou J, Chee MWL. 2016. Spontaneous eyelid closures link vigilance fluctuation with fMRI dynamic connectivity states. Proc Natl Acad Sci 113:9653–9658. doi:10.1073/pnas.1523980113

Whittaker JR, Driver ID, Venzi M, Bright MG, Murphy K. 2019. Cerebral autoregulation evidenced by synchronized low frequency oscillations in blood pressure and resting-state fMRI. Front Neurosci 13:1–12. doi:10.3389/fnins.2019.00433

Winder AT, Echagarruga C, Zhang Q, Drew PJ. 2017. Weak correlations between hemodynamic signals and ongoing neural activity during the resting state. Nat Neurosci 20:1761–1769. doi:10.1038/s41593-017-0007-y

Wise RG, Ide K, Poulin MJ, Tracey I. 2004. Resting fluctuations in arterial carbon dioxide induce significant low frequency variations in BOLD signal. Neuroimage 21:1652–1664. doi:10.1016/j.neuroimage.2003.11.025

Wong CK, Zotev V, Misaki M, Phillips R, Luo Q, Bodurka J. 2016. Automatic EEG-assisted retrospective motion correction for fMRI (aE-REMCOR). Neuroimage 129:133–147. doi:10.1016/j.neuroimage.2016.01.042

Wong CW, Olafsson V, Tal O, Liu TT. 2013. The amplitude of the resting-state fMRI global signal is related to EEG vigilance measures. Neuroimage 83:983–990. doi:10.1016/j.neuroimage.2013.07.057

Xia CH, Ma Z, Ciric R, Gu S, Betzel RF, Kaczkurkin AN, Calkins ME, Cook PA, García de la Garza A, Vandekar SN, Cui Z, Moore TM, Roalf DR, Ruparel K, Wolf DH, Davatzikos C, Gur RC, Gur RE, Shinohara RT, Bassett DS, Satterthwaite TD. 2018. Linked dimensions of psychopathology and connectivity in functional brain networks. Nat Commun 9:1–14. doi:10.1038/s41467-018-05317-y

Xia Y, Chen Q, Shi L, Li MZ, Gong W, Chen H, Qiu J. 2019. Tracking the dynamic functional connectivity structure of the human brain across the adult lifespan. Hum Brain Mapp 40:717–728. doi:10.1002/hbm.24385

Yan CG, Cheung B, Kelly C, Colcombe S, Craddock RC, Di Martino A, Li Q, Zuo XN, Castellanos FX, Milham MP. 2013. A comprehensive assessment of regional variation in the impact of head micromovements on functional connectomics. Neuroimage 76:183–201. doi:10.1016/j.neuroimage.2013.03.004

Zeng L-L, Wang D, Fox MD, Sabuncu M, Hu D, Ge M, Buckner RL, Liu H. 2014. Neurobiological basis of head motion in brain imaging. Proc Natl Acad Sci 111:6058–6062. doi:10.1073/pnas.1317424111

Zhang J, Huang Z, Tumati S, Northoff G. 2019. Intrinsic Architecture of Global Signal Topography and Its Modulation by Tasks. bioRxiv.

Zilles K, Armstrong E, Schleicher A, Kretschmann HJ. 1988. The human pattern of gyrification in the cerebral cortex. Anat Embryol (Berl) 179:173–179. doi:10.1007/BF00304699

